# Reassessment of the risks of climate change for terrestrial ecosystems

**DOI:** 10.1101/2023.10.24.563567

**Authors:** Timo Conradi, Urs Eggli, Holger Kreft, Andreas H. Schweiger, Patrick Weigelt, Steven I. Higgins

## Abstract

Forecasting the risks of climate change for species and ecosystems is necessary for developing targeted conservation strategies. Previous risk assessments mapped the exposure of the global land surface to changes in climate^1–4^ However, this procedure is unlikely to robustly identify priority areas for conservation actions because non-linear physiological responses and co-limitation processes ensure that ecological changes will not map perfectly to the forecast climatic changes. Here, we combine ecophysiological growth models of 135,153 vascular plant species and plant growth form information to transform ambient and future climatologies into phytoclimates, which describe the ability of climates to support the plant growth forms that characterise terrestrial ecosystems. We forecast that 33% to 68% of the global land surface will experience a significant change in phytoclimate by 2070 under RCP 2.6 and RCP 8.5, respectively. Novel phytoclimates without present-day analogue are forecast to emerge on 0.3-2.2% of the land surface, and 0.1-1.3% of currently realised phytoclimates are forecast to disappear. Notably, the geographic pattern of change, disappearance and novelty of phytoclimates differs markedly from the pattern of analogous trends in climates detected by previous studies^1,3,4^, thereby defining new priorities for conservation actions and highlighting the limits of using untransformed climate change exposure indices in ecological risk assessments. Our findings suggest that a profound transformation of the biosphere is underway and emphasise the need for a timely adaptation of biodiversity management practices.

## Main

Global Circulation Models forecast significant climatic changes throughout the 2l8t century under all but the most optimistic greenhouse gas emission scenarios^5^. The anticipated climatic changes are expected to force both continuous and abrupt changes in the distribution of ecosystems and species^6, 7^. One implication is that ecosystem managers may have to shift their focus from targeting pre-defined baseline states to managing ecosystem change along trajectories forced by climatic change8-^10^. However, the strength of climatic forcing and the direction of the new trajectories are uncertain, making it difficult for ecosystem managers to define and implement targeted actions^11, 12^. Not only are climates changing, but it is also likely that climate states without present-day analogues (hereafter novel climates) will emerge and that some of the existing climate states will disappear (hereafter disappearing climates)l, ^3^. It has been suggested that disappearing climates may increase the risk of loosing species and some types of ecosystems, whereas novel climates may lead to the formation of novel ecosystems^1, 13, 14^. Since the functioning of novel ecosystems is by definition unknown, their emergence would further enhance the risk of ecosystem management failure^1, 14, 15^. It is thus a research priority to identify regions where climate change is likely to force strong ecological change, so that ecosystem managers can implement timely and targeted actions.

To identify regions with elevated ecological risks from climate change, previous works analysed the exposure of the global land surface to potentially dangerous climatic changes^3^. This includes analyses of the exposure of ecosystems to strong climatic changes and globally novel climates, as well as the disappearance of existing climate states^1,4^. In these studies, exposure was calculated as the Euclidean distance between ambient and future climates, standardized by the historical variability in climate variables. Another widely used climate­ change exposure metric is climate change velocity^2, 16, 17^, which quantifies the displacement rate of climate states, and a more recent species-focused study identified where and when animal species will be exposed to mean temperature and precipitation conditions outside their realised niches^7^.

However, risk assessments based on climate-change exposure indices oversimplify an organism’s perception of climate-exposure. Previous climate-change exposure work did not consider that physiological and ecological responses to changing climate parameters are often non-linear and co-limited by multiple climate parameters^18–21^, and that hierarchies of co-limiting climatic parameters are expected to reorganise as climatic change progresses^22^•^23^. This means that the risk of a one unit increase in a climate parameter is not comparable across locations with different baseline parameter values and is contingent on concomitant changes in co-limiting climate parameters. Neither this baseline effect nor the co-limitation effect are accounted for by climate change exposure studies. Moreover, individual species and growth forms exhibit contrasting climatic preferences^24^ and are therefore likely to respond differently to projected climatic changes. The implication is that the ecological response to altered climatic conditions is unlikely to map perfectly to climate change exposure indices^1, 25^, which makes the ecological interpretation of climate-change exposure indices problematic.

Problems with the ecological interpretation of exposure indices could be addressed by using process-rich ecosystem simulation models^26, 27^. Such models are explicitly designed to model the impacts of climatic changes on ecosystems. However, ecosystem simulation models are hampered by process and parameter uncertainty^26–28^, evidenced by large discrepancies between outputs of different models forced with the same climate data^29^. The implication is that current approaches to understand ecological climate-change risks and impacts trade-off the certainty with which we can make predictions with the ability to ecologically interpret those predictions. Exposure indices have more tractable prediction uncertainty, yet are difficult to interpret ecologically, whereas the opposite is true for ecosystem models.

### Translating climates into phytoclimates

To reduce this trade off, we computed a phytoclimatic transform of ambient (mean of 1979-2013) and future (mean of 2061-2080; henceforth 2070) climatologies. The phytoclimatic transform expresses a grid cell’s climate in terms of its suitability for species of 14 plant growth forms that define terrestrial ecosystems. The transformation was based on an existing protocol^24^ (Fig. E1) and involved (1) parameterising an ecophysiological plant growth model forced with monthly climate data for 135,153 vascular plant species, (2) using the fitted species models to identify climatically suitable grid cells for each species, (3) calculating the proportion of species of each growth form for which a grid cell is climatically suitable and using this proportion as an index of a grid cell’s climatic suitability for a growth form (Fig. 1). We refer to a grid cell’s vector of the 14 growth form suitabilities as its phytoclimate. That is, rather than describing a location’s climate by climate variables such as mean annual temperature or annual rainfall, we describe the climate by its ability to support species of different types of plants that ecologists use to define terrestrial ecosystems. This provides a means to understand which structural changes in ecosystems the future climatic states will promote. For instance, should a grid cell’s climatic suitability for cold-deciduous trees change from 0.2 to 0.3, this would mean that hat the climate can now accommodate an additional 10% of species of this growth form at that location, which raises the potential for species of this growth form to become more frequent.

**Figure 1.**
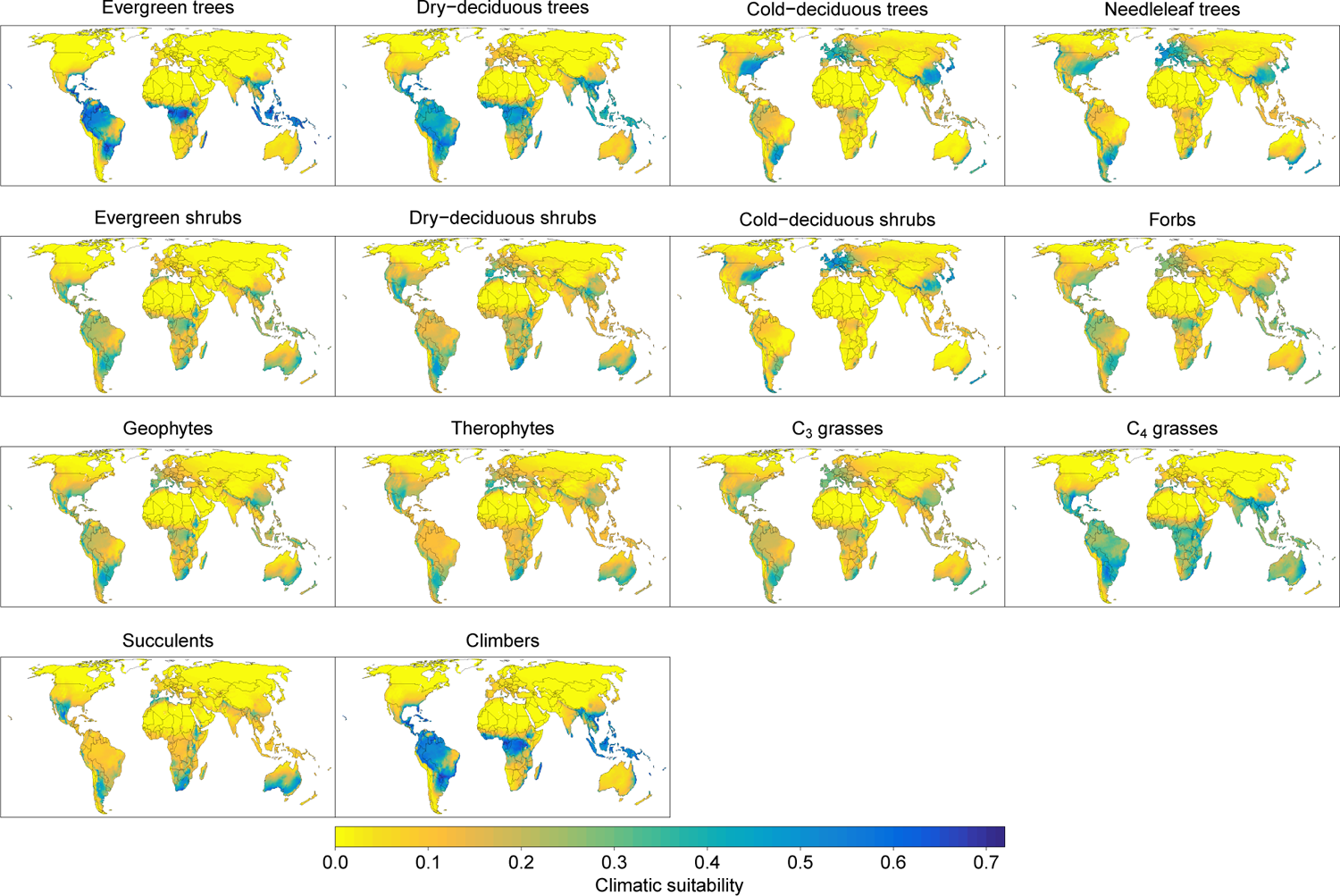
Ambient climatic suitability for major plant growth forms. Suitability is expressed as the proportion of plant species of a growth form for which the cells’ climate is suitable according to an ecophysiological plant growth model. The median number of modelled species per growth form was 5,249, with a minimum of 439 (needleleaf trees) and a maximum of 24,853 (evergreen shrubs) species. The total number of modelled species was 135,153.

The premise of this analysis is that we can characterize changes in the climatic forcing of ecosystems by analyzing shifts in climatic suitability for the growth forms that define ecosystems. Previous work showed that the growth-form suitability values are useful predictors of ecosystem states^30^, suggesting that shifts in growth form suitability may indeed be useful indicators of the changing climatic forcing. It should be emphasised that the growth-form suitability may be quite different from growth-form abundances, because other ecological processes such as biotic interactions, dispersal or disturbances determine whether physiological suitability can predict abundance. Our analysis therefore provides a physiologically-informed assessment of where and how the climatic drivers of ecosystem assembly are changing.

The plant growth model used here describes how the uptake and allocation of carbon and nitrogen of an individual plant is co-limited by monthly temperature, soil moisture, solar radiation and atmospheric CO2 concentrations^24,31,32^. We used species distribution data from the BIEN database (https://bien.nceas.ucsb.edu/bien/) to find the physiological parameters of the plant growth model that best explain the observed distribution of a species. For each of the 135,153 species, the parameterised model is then run forward using a monthly gridded climatology to simulate the biomass accumulation of the species in the cells of a global grid, and the simulated biomass values are used as the linear predictor in a logistic regression model that predicts whether grid cells are climatically suitable for that species or not. That is, we use data on the realised distribution of species to estimate the parameters of a model that articulates a simplified physiological niche of a plant species^32^. The performance of the model **in** transferability tests^33^ indicates that this physiological niche characterisation is predictive of where a plant species could grow.

The growth form suitability surfaces in Fig. 1 represent a summary of estimated physiological niches of 135,153 plant species grouped by growth form. We aggregate this phytoclimatic transform of the climate by identifying groups of cells with similar growth form suitability using unsupervised classification. The geographical projection of these groups reveals phytoclimatic zones of the Earth (Fig. 2A) and provides a plant growth-form centric classification of the Earth’s climates. Table E1 and Fig. E2 provide overviews of the mean growth form suitabilities of the ambient phytoclimatic zones. Figure E3 shows that the phytoclimatic zones have distinct climatic characteristics.

**Figure 2.**
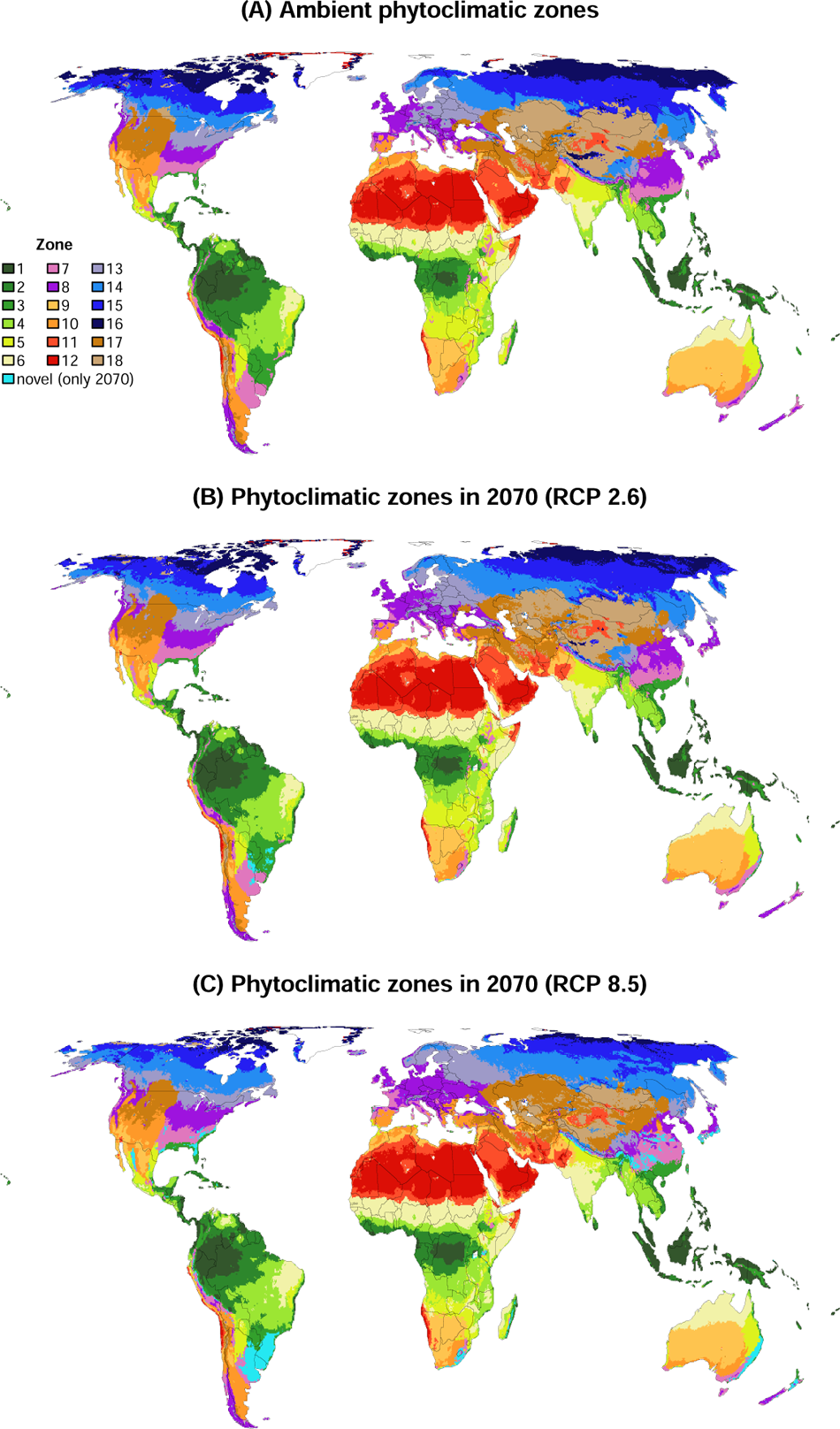
Phytoclimatic zones of the Earth and their shifts by 2070. Phytoclimatic zones have internally similar climatic suitability for different plant growth forms. (A) shows ambient phytoclimatic zones, derived from an unsupervised classification of grid cells by their climatic suitability for 14 plant growth forms (Fig. 1). (B) and (C) show the arrangement of phytoclimatic zones in 2070 under RCP 2.6 and RCP 8.5, respectively. The median climatic suitabilities for each growth form across five future climatologies were used to determine the future phytoclimatic zone of the grid cells for each RCP. Cells shown as ‘novel’ are projected to have growth form suitability realisations without present-day analogue. Figure E12 shows shifts in phytoclimatic zones for different combinations of RCP and GCM.

The phytoclimatic zones depicted in Fig. 2A represent areas where the climate supports similar plant types, making the concept closely related to biomes. Indeed, previous work suggested that phytoclimatic zones could be used to define biomes^24^. Yet, biomes are mostly defined as regions where specific combinations of plant growth forms have developed in response to the regional climate^34,35^. That is, biomes implicitly consider additional ecological processes that shape the ecosystem structure observed in the field, whereas phytoclimatic zones consider only how the climate influences the physiological performance of plant types. This means that phytoclimatic zones (e.g. Fig. 2) are more closely affiliated with bioclimatic zones^36,37^ than with biomes.

The phytoclimatic transformation was then applied to climatologies projected for 2070 under the RCP 2.6 and RCP 8.5 scenarios, which represent a reduced and a high emission scenario, respectively. Specifically, we forced the fitted species growth models with the future climatologies to project climatic suitability maps for each species and grouped species by growth form to calculate growth form suitability maps for 2070 (Figs. E4-E5).

Analogous to work on climate-change exposure^1^, we used the ambient and future growth form suitability maps to calculate three multivariate Euclidean distances that summarise components of ecological risk: (i) The local phytoclimatic change each grid cell will be exposed to, estimated as the distance between the ambient and future growth form suitabilities of each grid cell. This index summarizes the changing climatic constrains on the plant growth forms in each cell. (ii) The novelty of each grid cell’s future phytoclimatic state relative to ambient phytoclimatic states, estimated as the minimum distance between a grid cell’s future growth form suitabilities and that of an ambient grid cell. A novel phytoclimate is thus a climate that constrains plant growth forms differently to any ambient climate state. **(iii)** The disappearance of each grid cell’s ambient phytoclimatic state, estimated as the minimum distance between a grid cell’s ambient growth form suitabilities and that of a future grid cell. The disappearing phytoclimate index is thereby a measure of how distinct the ambient phytoclimatic state of a grid cell is relative to the future distribution of phytoclimatic states. A disappearing phytoclimate indicates disappearance of the way by which an ambient climates constrains plant growth forms.

To interpret which values of the three indices indicate high risks, we use the Euclidean distances between the phytoclimatic zones shown in Fig. 2A as a reference. Specifically, we computed the centroids of each zone in Euclidean growth form suitability space and calculated the pairwise Euclidean distances between these centroids. The 5^th^ percentile of these inter-centroid distances was used as a threshold value to identify risk equivalent to a shift between some of the phytoclimatic zones. Figures E6-E8 show the sensitivity of the results to varying the threshold value. Lastly, the future phytoclimatic zone of grid cells was projected by assigning grid cells to the zone of the closest ambient grid cell in Euclidean growth form suitability space; grid cells that were further than the threshold distance from ambient phytoclimatic states were designated as novel.

### Change, disappearance and novelty of phytoclimates

Our analysis predicts substantial change in phytoclimates by 2070 (Fig. 3A). If anthropogenic emissions follow RCP 2.6, 33% of the Earth’s terrestrial surface (excluding the currently ice­ covered parts of Greenland and Antarctica) will experience an ecologically significant change in the extent to which the climate can support different plant growth forms. The fraction of land with significant change in phytoclimate increases to 68% if emissions follow RCP 8.5. When calculating the median change value across five projections that used climatologies from different Global circulation Models (GCMs), the most pronounced phytoclimatic changes are likely to occur in the mountain regions of south China, the Himalayas, northwestern Russia, the Baltic countries, Scandinavia, the south- and northeastern United States, Alaska, central Mexico, the tropical Andes, southeastern South America, southeastern Australia and northern New Zealand. Projections based on each GCM are shown in Figs. E9A & E10A.

**Figure 3.**
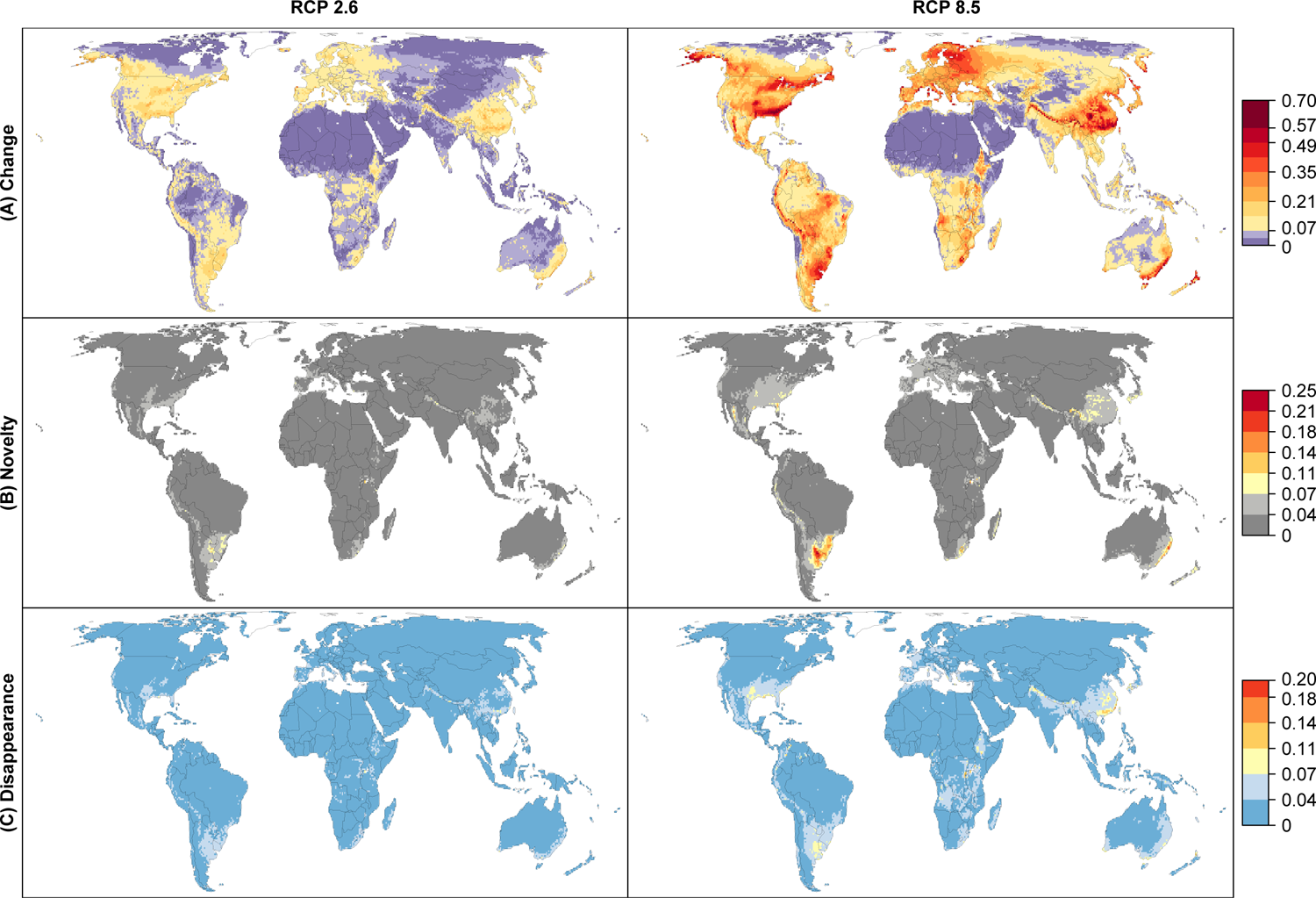
(A) Change, (B) novelty and (C) disappearance of phytoclimates by 2070 under RCP 2.6 (left column) and **8.5 (right column).** The phytoclimate of the cells is the suitability of the local climate for 14 plant growth forms that characterise the structure of terrestrial ecosystems. (A) Local change **in** phytoclimate, expressed as the Euclidean distance between a cell’s ambient and future phytoclimate. (B) Risk of novelty of the projected phytoclimate in 2070, expressed as the Euclidean distance of a cell’s future phytoclimate to its closest analogue in the global pool of ambient phytoclimates. (C) Risk of disappearance of the existing phytoclimate, expressed as the Euclidean distance of a cell’s ambient phytoclimate to its closest analogue in the global pool of future phytoclimates. In (A), (B) and (C), the colour bar is scaled so that yellow to red colours indicate significant values of change, novelty and disappearance, respectively (see main text for the definition of the threshold value) (Fig. 2A). Values are medians across future climatologies generated by five different GCMs for each RCP.

The changes in local phytoclimates translate into widespread shifts of phytoclimatic zones (Fig. 2B-C). Temperate, boreal and polar regions are most strongly affected by these shifts (Fig. E11). Under RCP 8.5, large parts of zones 13 and 14, which currently support cool-temperate and hemi-boreal ecosystems, will shift polewards, as will zone 15, which currently supports boreal ecosystems. Zone 16, which currently supports tundra ecosystems, will be reduced by 72% as it cannot shift further north. This prediction is consistent with observations of more frequent recruitment of shrubs and trees in tundra^38–41^ and increasing tree density at the forest-tundra ecotone^42^. Zone 18, which supports ecosystems of very continental cool-temperate regions will loose 54% of its current extent, mostly to zone 17, which favors less cold-limited continental cool-temperate ecosystems (Fig. E3) and is generally more suitable for most growth forms (Fig. E2). Tropical phytoclimatic zones are overall less likely to change in position and extent. However, some regions will be exposed to changes in phytoclimatic zones that may force strong structural changes in ecosystems. For instance, the southeastern and eastern parts of the Amazon shift to a phytoclimatic zone which supports savanna and some types of drier forest ecosystems (phytoclimatic zone 4, Fig. Ell). This projection is consistent with analyses that have used hydrological thresholds to project future changes in South American forests and savannas^43^. Phytoclimatic zone 4 is predicted to advance into the Amazon irrespective of GCM, albeit to very different degrees (Fig. E12), ensuring that the future phytoclimatic status of the Amazon region is highly uncertain (Fig. E13).

Despite the strong changes in local phytoclimates and substantial shifts in phytoclimatic zones, most of the future phytoclimates will have a present-day analogue. Specifically, only 0.3% (RCP 2.6) to 2.2% (RCP 8.5) of land cells will exhibit novel phytoclimates (Fig. 2B,C). The highest novelty indices are forecast in southeastern South America and Australia (Fig. 3B). The novel phytoclimates are forecast to primarily emerge in what at present are mesic subtropical climates typical of the eastern sides of continents, but they are also likely in tropical and subtropical mountain ranges such as the Andes, the Himalayas and the Sierra Madre Occidental. The majority of grid cells in the South American and Australian novelty centers belong to the ambient phytoclimatic zone 7, which is charaterised by relatively high suitability for all growth forms (Fig. E2). These cells are predicted to exhibit a strongly increased climatic suitability for evergreen trees, dry-deciduous trees and climbers by 2070, but reduced suitability for needleleaf trees, cold-deciduous shrubs and cold-deciduous trees (Figs. E14-E15). The emerging phytoclimatic novelty in these cells may be induced by slightly increasing rainfall combined with similar temperature seasonality, albeit with hotter summers and milder winters. It is likely that this combination reduces the cold-limitation of the numerous species of woody growth forms with preference for mesic tropical and subtropical climates in our analysis, whilst the elevated temperatures disfavor species of woody growth forms with preference for temperate conditions (Fig. 1).

Previous analyses of climatic variables found that the highest climatic novelty will emerge at the hot border of the ambient climate space in the tropics1, ^3^. Ouranalyses of phytoclimates suggests these novel tropical climates are not functionally novel. For instance, although parts of the Sahara will be exposed to novel climates^1^•^3^, we predict that these novel climates will remain highly unsuitable for all plant growth forms, which means that the novel climates are not novel from a plant functional perspective. In the wet tropics where decreases in rainfall and higher temperatures are projected, the novel climates may promote more drought adapted species, yielding growth form suitability combinations (phytoclimates) already observed elsewhere. For example, the Amazon and parts of the Cerrado are novel climate regions^1^•^3^, yet we predict that phytoclimates currently realised in the Cerrado are advancing in the Amazon, and are themselves partially replaced by phytoclimates currently realised in the Caatinga (Fig. 2C), meaning that no novel phytoclimates emerge in this region.

Only 0.1% (RPC 2.6) to 1.3% (RPC 8.5) of ambient phytoclimates are projected to disappear by 2070 (Fig. 3D). As with novelty, the disappearance of phytoclimates is forecast to occur primarily in the mesic subtropical climates on the eastern sides of the continents and there was good agreement between the five projections per RCP on where novel and disappearing phytoclimates are located (Figs. E9B,C & ElOB,C). Dissapearing phytoclimates were seldom in the same grid cells that were predicted to support novel phytoclimates. For instance, using the threshold applied in Fig. 3, only 25% of the grid cells with a disappearing phytoclimate were projected to be operating under a novel phytoclimate in 2070, whereas 75% of the cells with disappearing phytoclimates will be operating under a phytoclimate that is found elsewhere today.

The spatial resolution of this analysis can hide details of the geography of changing phytoclimates. While we fitted the species growth models using climatic forcing data with 1-km spatial resolution, our global analysis of phytoclimates used a grid with circa 25-km spatial resolution. This aggregation removes the (phyto-)climatic heterogeneity within the 25-km cells, preventing the detection of finer phytoclimatic patterns in topographic heterogeneous areas such as mountain ranges. For example the alpine tundra of the European Alps is not visible in our map of phytoclimatic zones (Fig. 2A). A higher resolution would allow the presence and dynamics of such micro-phytoclimates to be projected.

Moreover, while we explored uncertainty originating from RCP and GCM (Figs. E9, E10, E12, E13), we did not explore uncertainty originating from the model used to estimate climatic suitability for the individual plant species. Our growth form suitability projections are therefore partly contingent on the assumptions of the growth model used to estimate species-level suitability. While in principle any suitability or species distribution model could be used in the workflow, in practice, many existing alternative models may be inappropriate for this task. First, correlative suitability models have limited transferability^33^•^44^, producing implausible growth form suitability surfaces^24^, whereas the process-based TTR-SDM used here was successful in model transferability tests^33^ and produces plausible suitability surfaces^24^•^30^.

Second, alternative models do not consider how elevated atmospheric CO2 concentrations may influence photosynthesis and species ranges. Third, sensitivity analyses (Fig. E19) show that robust estimates of growth form suitability surfaces require on the order of hundreds of species distribution models, meaning that only process-based models that can be calibrated for many species are suitable for this task. On another level, uncertainty in the species distribution model needs to be evaluated in the context of other significant forms of uncertainty, such as uncertainty in the occurrence data, the assignment of species to growth form classes, and the climatic forcing data used to calibrate the models and to project them into the future.

### Implications of future phytoclimates

Global greenhouse gas emissions are currently tracking the RCP 8.5 scenario^45^. Under this scenario, our analysis predicts that almost the entire vegetated land surface will be subject to substantial changes in how climate supports the different types of plants that define terrestrial ecosystems (Fig. 3A). This is likely to impact on ecosystem structure, functioning and dynamics. For example, successional trajectories may no longer follow the usual sequences^21,46^ and some undesired growth forms may increase in abundance^47^, complicating ecosystem restoration and management. The prospect of changes in ecosystem structure and functioning supports growing calls for ecosystem managers to realign their goals and practices^8–10^. For instance, conservation management in protected areas often seeks to retard change by removing invading species, but if climatic forcing threatens the baseline states, actions that enhance ecosystems’ ability to track climatic forcing may be more effective^9,10^. Tracking climatic forcing would require that managers plan actions that ensure that locally declining species can reach suitable locations elsewhere, for instance species translocations^48^ or enhancing the permeability of agricultural landscapes^49^. Management actions that aim to retard change need to be carefully planned since such actions, if not sustained, will be a waste of resources^50^. Similarly, in ecological restoration it is common to use pre-defined baseline states as target points^51^, but these targets will become increasingly unattainable as climate deviates from domains that allow the targeted baseline states^52^. Our analysis can be used as a guide to identify locations where future climates will not support current ecosystem states and where managers could consider switching from preservation to managing the trajectories of change^9,10^. Our projections of shifts in phytoclimatic zones (Fig. 2) may serve to reduce uncertainty about these trajectories, because they show which type of phytoclimate is expected in a locale in the future and where experience with managing ecosystems under such a phytoclimate may already exist.

Our analysis predicts how the climatic forcing of ecosystems is changing, but other factors, not considered in our analysis, will influence the extent and rate at which ecosystems follow the trajectories of climatic forcing. In particular, disturbances or dieback of late-successional vegetation can accelerate change in ecosystems^21^. Other factors, such as dispersal limitation^53^, priority effects of resident communities^54^, persistence of declining remnant populations^55^ or microclimatic buffering by tree canopies^56^ can cause ecosystems to lag behind the climatic forcing, leading to a disequilibrium between climate forcing and ecosystem state. An ecosystem in strong disequilibrium with climate is however at high risk of sudden and potentially undesired transitions when the barriers to change have been overcome^32^, underlining that it may be prudent to proactively manage the change in ecosystems in order to reduce risks associated with mismatches between climate forcing and ecosystem states. This could be achieved by various means including assisted migration, increasing landscape connectivity^48,49,57^ and rewilding^58,59^. All of these require authentic shifts in management paradigms, which may be warranted given that the majority of the land surface will, according to our analysis, be subject to fundamental changes in climatic forcing.

Novel phytoclimates also have implications for ecosystem management. Novelty in our analysis emerges when the climate supports plant growth forms in unprecedented ways. The emergence of novel phytoclimates confronts biodiversity managers with deeper uncertainty on how ecosystems respond to management actions^15^ and how the ecosystems might function and look structurally in the future. That is, where novel phytoclimates emerge, managers cannot rely on experience gained by managers elsewhere. The palynological record indicates that no-analogue climates in the last late-glacial supported vegetation formations without a present-day analogue^60^, suggesting that the novel phytoclimates of the future may, as they have done in the past, give rise to novel ecosystems.

The interpretation of disappearing phytoclimates is that some of the ways in which climate currently supports plant growth forms will not be realised in the future. As these phytoclimates disappear, there is potential for the ecosystem types that form under these phytoclimates to disappear. It follows that regions in which we project disappearing phytoclimates are high risk areas for biodiversity loss because it is unlikely that these ecosystem states can assemble or be restored elsewhere. We found a coherent pattern in the distribution of disappearing and novel phytoclimates, suggesting priority areas for conservation monitoring and action.

The hotspots of phytoclimatic change, novel phytoclimate emergence and phytoclimate disappearance predicted in this study (Fig. 3) differ clearly from regions of analogous climatic changes identified in previous global studies^1^•^3^. These studies used climate system variables of broad ecological relevance that were standardised against their inter-annual variability in the time period covered by the ambient climatology. This standardisation emphasises changes in variables that are large relative to their historical variability, because such changes are assumed to have stronger ecological effects, and tends to up-weight changes in temperature over changes in precipitation since the inter-annual variability of temperature is often smaller^1^. Our approach by contrast uses an ecophysiological growth model to transform monthly climate variables into estimates of climatic suitability for individual species and, by grouping species into growth forms, the climatic suitability for the plant growth forms defining terrestrial ecosystems. That is, the ecological effect of temporal changes in climate variables is prescribed by the fitted plant growth models and summarised in the future growth form suitability surfaces. This phytoclimatic transformation explains why tropical regions, which are predicted to produce novel climates due to warmer temperatures^1^•^3^, are not predicted to produce novel phytoclimates as warming proceeds. Specifically, as warming proceeds, the suitability for all growth forms is forecast to decline in the equatorial regions (Figs. E14-E15) and the phytoclimate in these regions will become similar to phytoclimatic zones supporting more drought-adapted and seasonal ecosystems that already exist elsewhere (Fig. 2B,C).

Global greenhouse gas emissions are currently tracking RCP 8.5 predictions^45^ under which severe climatic changes are forecast^5^. Our analysis suggests that these climatic changes are likely to force a profound transformation of the biosphere, ensuring that substantial adaptation measures will be necessary for biodiversity conservation, agriculture and forestry. Our findings however also show that changes to the biosphere would be considerably milder under RCP 2.6, which supports the view that cutting greenhouse gas emissions would fundamentally reduce climate change risks for biodiversity, ecosystem functioning and agricultural production.

## Methods

### Environmental data

The plant growth model (described below) is forced with data on monthly minimum, mean and maximum temperature, monthly soil moisture, monthly solar radiation, and atmospheric CO2 concentrations. Nitrogen uptake is simulated as a function of temperature, soil moisture and soil nitrogen content^32^. In this application of the model we assumed the same soil nitrogen content in all grid cells, which means that plant nitrogen uptake in the model is influenced by temperature and soil moisture only. This is appropriate because this analysis aims to estimate the climatic suitability of geolocations. Ambient monthly temperature data (averages for the period 1979-2013) were downloaded from the CHELSA database^61^ version 1.2. Solar radiation data were downloaded from the Global Aridity and PET Database^62^. We developed a soil moisture model that is similar to that of ref^62^ and predicts monthly soil moisture based on monthly values of precipitation, solar radiation, minimum, mean and maximum temperature, and soil field capacity and wilting point, using a Hargreaves-type model of monthly potential evapotranspiration. Ambient monthly precipitation data (averages for the years 1979-2013) were taken from CHELSA version 1.2, and soil field capacity and wilting point data from the International Geosphere-Biosphere Programme Data and Information System^63^. The modelled soil moisture values reflect soil moisture available for evapotranspiration and are not influenced by the vegetation of a grid cell. For model fitting, we assumed an ambient atmospheric CO2 concentration of 338 ppm^64^. All environmental data were resampled to 1-km resolution if necessary and projected to the equal area World Eckert IV projection, which was used in all analyses and maps.

We downloaded ten downscaled CMIP5 temperature and precipitation climatologies for 2070 (averages for the period 2061-2080) projected by five CMIP5 Global Circulation Models (GCMs) under two emission scenarios (RCP 2.6 and 8.5) from CHELSA version 1.2. The five GCMs were: CCSM4^65^, CNRM-CM5^66^, FGOALS-g2^67^, MIROC-ESM^68^, MPI-ESM­LR^69^. These models were chosen to represent a wide range of uncertainty in climate-change projections originating from different GCMs^70^. Future monthly soil moisture was predicted with our soil moisture model for each of the ten climatologies for 2070. Solar radiation in 2070 was assumed to be the same as today. We focus on RCP 2.6 and 8.5 because the former represents an optimistic emissions reduction scenario whereas the latter represents a pessimistic scenario that assumes continued growth in emissions^71^ and is the scenario the world is currently tracking most closely^45^. We assumed atmospheric CO2 concentrations in 2070 of 438 ppm and 677 ppm for RCP 2.6 and 8.5, respectively^64^.

We classified the grid cells of a global map into 20 environmental zones based on the ambient monthly environmental forcing data used by the plant growth model (Fig. E16). We used the *clara* algorithm in the *cluster* package^72^ in R to classify the cells and optimised the separation of cells into the 20 *clara* clusters by means of a Discriminant Analysis of Principal Components (DAPC)^73^ computed from the environmental data. The resulting environmental zones were used later for generating stratified samples of species presence and pseudo-absence records that were used to fit the plant growth model.

### Species distribution and growth form data

We downloaded distribution data of all vascular plant species in the BIEN database version 4.1 (http://bien.nceas.ucsb.edu/bien/), using the BIEN R package^74^. We used the non-public version of BIEN, which contains sensitive occurrence data of endangered species not included in the public BIEN version and was made available to us upon request.

Most occurrence records in BIEN come from herbarium collections (see acknowledgment section for collections used in this analysis), ecological plots and surveys7, ^75^ as well as from plant trait observations. BIEN also includes data from NeoTropTree (http://www.neotroptree.info/), RAINBIO (http://rainbio.cesab.org/), TEAM (https://www.wildlifeinsights.org/team-network) and The Royal Botanical Garden of Syd­ ney, Australia (https://www.rbgsyd.nsw.gov.au/) and plot data from CVS, CTFS, FIA, NVS, SALVIAS, TEAM, VEGBANK and MADIDI (see https://bien.nceas.ucsb.edu/bien/data-contributors/all/). A full list of references for BIEN occurrence records used in this study can be found in Extended Data Table E2.

The R package CoordinateCleaner version 2.0-11^76^ was used to remove records with either zero longitude or latitude, and records within a buffer of 10 km around country and province centroids, 5 km around country capitals, and 200 m around biodiversity institutions (herbariums, museums or universities), respectively. In addition, we computed country centroids from the Database of Global Administrative Areas version 3.4 and removed records within a 50-km buffer to these centroids. We only retained one occurrence record per 1-km grid cell and species.

The species were grouped into 14 growth forms: evergreen broadleaf trees and shrubs, cold­ and drought-deciduous broadleaf trees and shrubs, needleleaf trees, C3 and C4 grasses, forbs (excluding geophytes and therophytes), geophytes, therophytes, terrestrial succulents, and climbers. All Pinales except Gnetales and Podocarpaceae were classified as needleleaf trees, that is all Araucariaceae, Cephalotaxaceae, Cupressaceae, Pinaceae, Sciadopityaceae and Taxaceae. All species of the Poales families Poaceae, Cyperaceae, Juncaceae, Anarthriaceae, Centrolepidaceae, Ecdeiocoleaceae, Joinvilleaceae, Restionaceae, Thurniaceae and Typhaceae were classified as grasses. The database of ref^77^ was then used to identify grass species with C4 photosynthetic pathway. We classified all species listed in the Illustrated Handbook of Succulent Plants as succulents^78^, with updated lists for monocotyledons^79^ and Cactaceae^80^. Woody succulents were treated as succulents and not as trees or shrubs.

For all remaining species we extracted information on growth form and leaf phenology from BIEN and GIFT version 2.1^81^. We first searched for species-level information in BIEN and filled the gaps with species-level information in GIFT. If growth form and phenology data was still missing for a species, we searched for genus-level information in BIEN and filled the gaps with genus-level information in GIFT. If data was still missing after this step, we searched for family-level information in BIEN and filled the gaps with family-level information in GIFT. If multiple entries with contrasting information were available (e.g. a species had entries as a shrub and a tree), we used the most frequent entry. We assigned species classified in BIEN and GIFT as herb or fern to the forb category. Species classified as graminoids by BIEN and GIFT were also classified as forbs by us, unless they belonged to one of the Poales mentioned above. The forbs were then split into geophytes, therophytes and other forbs using life form information in GIFT. We assigned species classified in BIEN as climber, liana or vine to the climber category, as well as species classified in GIFT as obligatory climbers, liana or vine. Palms were treated as trees. We excluded tree and shrub species with leaf phenology entries “variable” (GIFT) or “variable or conflicting information” (BIEN), aquatic species and all epiphytes (incl. succulent epiphytes) because they do not use soil moisture and thus cannot be modelled with our approach (GIFT includes a comprehensive global checklist of vascular epiphytes^82^ that was used to identify epiphytes). Cold- and drought deciduousness of trees and shrubs was determined by plotting the BIEN distribution records on a world map of Koppen-Geiger climates^83^. Species with >50% of records in cold climates (Koppen-Geiger zones Cf, D and E) were defined as cold-deciduous and the remaining species were assumed to be drought-deciduous. References for BIEN growth-form data can be found in Extended Data Tables E3.

Name matching between data sources was accomplished with the Taxonomic Name Resolution Service^84^, which uses the Missouri Botanical Garden’s Tropicos database (https://tropicos.org), The Plant List (http://www.theplantlist.org) and the USDA Plants Database^85^.

### Growth and species distribution modelling

We used the TTR-SDM^32^, an ecophysiological species distribution model for plants, to identify climatically suitable grid cells for each plant species. The TTR-SDM is based on Thornley’s Transport Resistance (TTR) model^31^, which describes in a series of dynamic equations how the biomass accumulation of an individual plant is influenced by the uptake of carbon and nitrogen, their allocation between sinks and sources, and growth processes. The TTR-SDM^32^ includes a series of functions that describe how the resource uptake (carbon and nitrogen) and growth processes in the TTR model are influenced by monthly minimum, mean and maximum temperature, soil moisture, solar radiation, soil nitrogen and atmospheric CO2 concentration. Figure E20 provides a graphical representation of how these environmental forcing variables influence the resource uptake and growth rates in the TTR-SDM. The model’s equations prescribe the general shape of these relationships (i.e. trapezoidal or saturating), however the values of the forcing variables at which the physiological rates change are species-specific and are estimated for each plant species separately; these values are the model’s parameters (n=18; see Fig. E20) and we estimate them from species distribution data as described below. The model’s equations define that each of the forcing variables can co-limit a plant’s resource uptake and growth analogous to Liebig’s law of the minimum, and that allocation of the assimilated resources between resource sources and sinks is driven by transport resistance processes (see ref^32^). The model is run on a monthly time step using the ambient and the 2070 monthly climatology, respectively, which allows it to explicitly simulate how monthly fluctuations in the forcing variables co-limit a plant’s resource uptake, allocation and growth. That is, the model simulates that a plant’s monthly biomass accumulation is co-limited by the temperature, soil moisture, solar radiation, and soil nutrients that a plant is exposed to each month of a simulation^33^.

The TTR-SDM version used here uses a Farquhar-type photosynthesis model^86^ to describe how potential carbon assimilation rates are co-limited by light, temperature and atmospheric CO2 concentration and how this co-limitation differs for C3 and C4 plants (see details in ref^24^). We assume that each species uses either the C3 or the C4 photosynthetic pathway and use universal parameterisations of the C3 and C4 Farquhar models. Therefore, each grid cell has a universal maximum rate of monthly C3 or C4 carbon uptake that is determined by light, temperature and atmospheric CO2, and this universal maximum rate can be further reduced by species-specific shoot-nitrogen and soil-water dependencies of carbon uptake (Fig. E20).

The prescribed way in which environmental factors influence physiological processes (Fig. E20), the simulation of monthly co-limitation dynamics and the explicit consideration of CO_2_ effects via a Farquhar-type carbon assimilation model are key conceptual differences to correlative species distribution models. These properties allow the model to extrapolate in physiologically plausible ways to novel data domains, thereby accommodating both novel data ranges and novel combinations of monthly values of climate variables. For example, a model comparison^33^•^87^ between the TTR-SDM and the widely used correlative SDM Maxent^88^ showed that although the TTR-SDM has slightly lower ability to describe the species distribution data in the climate-data domain used to fit the models, it had a substantially better ability to describe independent species distribution data outside the climate-data domain used to fit the models. This model comparison provides confidence that the TTR­ SDM accurately identifies suitable climatic conditions.

To parameterise the model, one could measure the 18 model parameters shown in Fig. E20 in the laboratory, but this is not feasible when the goal is to parameterise the model for many species. The alternative is to infer the parameters from species distribution data and gridded climatologies. Conceptually, we achieve this as follows: First, we use the monthly climatic forcing data to simulate the biomass growth of a species at its presence and absence locations. Once the simulated biomass reaches equilibrium with the climate system forcing data, we use the natural log of this simulated biomass as the linear predictor in a logistic regression model that predicts the observed presences and absences of the species. The simulated biomass values are skewed and in such cases the complementary log-log link function is recommended. We therefore use the complementary log-log link function in this study. In practice, using logit or the complementary log-log often does not make a large difference^89^, even if there are theoretical reasons to prefer the complementary log-log. The inference process then uses an optimization algorithm to iteratively improve the likelihood of this regression model by optimising the **18** parameters of the growth model (Fig. E20), which are constrained by the prescribed trapezoidal and step functions defined by the model’s equations^32^. This optimization was performed using the Differential Evolution genetic algorithm^90^, a stochastic optimisation method implemented in R in the DEoptim package^91,92^. We allowed the algorithm to iterate 1000 times, which we found to be sufficient for generating stable parameterisations of the TTR-SDM.

We attempted to fit the TTR-SDM for species with at least seven presence data points. All presence points were used if less than 400 points were available. If more presence points were available, we took a random sample of presence points that conserved the proportions of the 20 environmental zones defined above (Fig. E16) in the full set of presence points. For species with more presence records than environmental zones, we then sampled the same number of pseudo-absence points as presence points (the actual number varied slightly due to integer rounding). The probability of selecting a pseudo-absence point in an environmental zone was inversely proportional to the proportions of the zones in the presence point sample, which ensured that zones strongly represented in the presence point sample were less likely to be included in the pseudo-absence points sample. For species with less presence records than environmental zones, we used 20 pseudo-absence points to better constrain the parameter estimation. We found in a pilot study using the benchmarking dataset from ref^33^ that our sampling strategy for pseudo-absence points produced parameterisations of the TTR-SDM that had the highest ability to predict independent species distribution data. Other tested sampling strategies used the same stratified strategy as above, but with 2, 5 and 10 times the number of pseudo-absence points as presence points, stratified sampling of pseudo-absence points without down-weighting, random sampling of pseudo-absence points and the target-group approach^93^, each with 2, 5 and 10 times the number of pseudo-absence points as presence points. These alternative strategies were found to generate parameterisations of the TTR-SDM with marginally lower predictive accuracy.

Once the final parameterisation is found, we use it to simulate a species’ potential biomass in the cells of a global grid using their monthly climatologies. The complementary log-log of the natural log of biomass is then used to calculate a suitability score (0-1). Lastly, to convert this suitability score into a binary prediction (0,1), we chose a threshold suitability score that maximises the sum of true positives and true negatives in the presence and (pseudo-)absence data used to parameterise the model. The result is a map showing where the climate is suitable for that species.

The predictive accuracy of the model is then evaluated using the True Skill Statistic (TSS) calculated from a confusion matrix^94^ that was computed using the above-mentioned threshold. Models with low predictive accuracy (TSS ≤0.7) were removed. This resulted in fitted models for 135,153 species, consisting of 24,362 evergreen trees, 3,173 dry-deciduous trees, 1,270 cold-deciduous trees, 439 needleleaf trees, 24,853 evergreen shrubs, 2,074 dry-deciduous shrubs, 1,943 cold-deciduous shrubs, 21,903 therophytes, 8,297 geophytes, 23,538 forbs, 6,888 C3 grasses, 2,814 C4 grasses, 3,609 succulents, 9,990 climbers.

The parameters of the species models were estimated using environmental data interpolated to a 1 km grid (see data sources). For subsequent data analyses these species models were projected onto a 25 km grid.

DEoptim is a robust and efficient global optimisation algorithm capable of finding optima on irregularly shaped likelihood surfaces^90^. The stochastic nature of the algorithm means that running DEoptim several times for the same species can produce different parameter estimates. This parameter uncertainty produces uncertainty about the potential ranges of individual species, which we use to calculate the proportion of species of each growth form that could grow in the grid cells (i.e. the growth form suitability values shown Fig. 1). The large number of species used in this analysis however ensures that the described species-level uncertainty is a negligible source of uncertainty about the growth form suitability values.

Figure E18 shows that taking five random samples of 50% of the species of a growth form and calculating the suitability scores of the grid cells from each sample yields almost identical suitability scores. In most growth forms, even smaller subsets would produce identical results (Fig. E19). The saturating curves in Figure E19 also shows that the number of species used in this study were sufficient to characterise the climatic preferences of each growth form. An additional sensitivity analysis showed that the estimation of the ambient suitability surfaces was robust to excluding all species with less than 20 occurrence records (Fig. El 7).

Biases in the species distribution data and the use of modeled coarse (1-km resolution) climate forcing data may bias the parameterization of the TTR-SDM. Model comparisons however show that the TTR-SDM is less biased than correlative SDMs in predicting locations where the climate is suitable for a species^33, 87^. Moreover, so long as these biases are not systematic, our procedure of averaging over multiple species models creates robust estimates of the growth form suitability surfaces (Figs. E17, E18, E19).

### Data analysis

For each 25-km grid cell, we calculated the proportion of species of each growth form that can grow in the cell according to the fitted TTR-SDM. This proportion is interpreted as the climatic suitability of a grid cell for a growth form (Fig. 1).

To find groups of cells with similar suitabilities for different growth forms we used finite Gaussian mixture modelling as implemented in the R package mclust^95^. We refer to the geographic projection of these groups as phytoclimatic zones (Fig. 2A). We estimated the Bayesian Information Criterion (BIC) for different variations of the mclust algorithm, which revealed that the option “VEV”, allowing ellipsoidal, equally shaped clusters, consistently performed better than the alternatives. We thus used this option to model the clusters.

Using the number of (terrestrial) biomes delimited by global biome maps to guide the optimal number of clusters, one might delimit for instance 13^96^, 14^97^, 20^98,99^, 21^100^ or 30^101^ clusters. We used 18 clusters to trade off information content versus interpretability. The BIC of the VEV clustering models improved with the number of clusters used up to 30 clusters, but indicated that the BIC of 18 clusters was not substantially lower than the maximum BIC at 30 clusters.

Our indices of novelty, disappearance and local change of phytoclimates, as well as the projections of shifts in phytoclimatic zones by 2070, are based on the multivariate Euclidean distances (ED) between ambient and future climatic suitabilities:

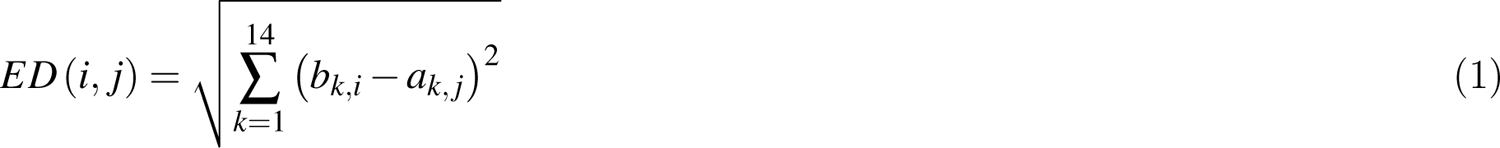

where ak_’_i and b k J. are the ambient and future suitabilities for growth form k in grid cell i and j, respectively. For within grid cell phytoclimatic change, i equals j. Note that the growth form suitabilitiy values all range from 0 to 1. We calculated the novelty of the future phytoclimate in grid cell x by setting i=x and calculating ED(i,j) for j=1 to N, where N is the total number of grid cells in the global 25 km grid. The minimum of these EDs is the distance of a future phytoclimate to its closest ambient analogue (EDmin)-A high EDmin thus indicates a high degree of novelty. Analogously, to determine phytoclimatic disappearance, we set j=x and calculate ED(i,j) for i=1 to N and calculate the minimum of these EDs. Here, high values of EDmin indicate that an ambient phytoclimate has no close future analogue. Figures E9 and ElO show phytoclimatic change, novelty and loss for each GCM separately. Figure 3 shows the median values across the five projections per RCP.

To generate projections of shifts in phytoclimatic zones by 2070, we calculated the minimum ED of a future phytoclimate to its closest ambient analogue and extracted the ambient phytoclimatic zone of this closest ambient analogue. The future phytoclimate was then assigned to this phytoclimatic zone. Figure E12 shows shifts in phytoclimatic zones for each combination of RCP and GCM. Figure 2B in the main text shows the projection of zone shifts based on the median growth form suitability values across the five projections for RC 2.6 and Figure 2C shows the same for RCP 8.5.

## Acknowledgments

We acknowledge the herbaria that contributed data to this work: A, AAH, AAS, AAU, ABH, ACAD, ACOR, AD, ADW, AFS, AHUC, AIMS, AJOU, AK, AKPM, ALCB, ALT, ALTA, ALU, AMD, AMES, AMNH, AMO, ANA, ANGU, ANSM, ANSP, ANUC, ARAN, ARC, ARIZ, ARM, AS, ASDM, ASU, ATCC, AUG, AUT, B, BA, BAA, BAB, BACP, BAF, BAFC, BAI, BAJ, BAL, BARC, BAS, BBB, BBS, BC, BCF, BCMEX, BCN, BCRU, BEREA, BG, BH, BHCB, BHO, BILAS, BIO, BISH, BLA, BM, BO, BOCH, BOG, BOL, BOLV, BONN, BOUM, BPI, BR, BRA, BREM, BRI, BRIT, BRIU, BRLU, BRM, BSB, BSIP, BSN, BTN, BUL, BULU, BUT, C, CALI, CAMU, CAN, CANB, CANL, CAS, CAY, CBG, CBM, CBS, CEN, CEPEC, CESJ, CGE, CHAM, CHAP, CHAPA, CHI, CHL, CHR, CHRB, CIB, CICY, CIIDIR, CIMI, CINC, CIQR, CLEMS, CLF, CM, CMC, CMMEX, CNHM, CNPO, CO, COA,COAH,COCA,CODAGEM,COFC,COL,COLO,CONC,CORD,CP,CPAP,CPUN, CR, CRAI, CRP, CS, CSU, CTES, CTESN, CU, CUVC, CUZ, CVRD, CWU, DAO, DAOM, DAV, DBN, DES, DLF, DMNH, DMU, DNA, DR, DS, DUKE, DUSS, E, EA, EAC, EBH, EBUM, ECH, ECU, EIF, EIU, EKY, EM, EMMA, ENCB, ENS, ERA, ESA, ESS, F, FAA, FAU, FB, FBCS, FCME, FCO, FCQ, FEN, FH, FHO, FI, FLAS, FLOR, FM, FR, FRP, FTG, FUEL, FURB, G, GB, GDA, GDAC, GE, GENT, GEO, GES, GH, GI, GJO, GLM, GMNHJ, GOET,GUA,GZU,H,HA,HAC,HAJB,HAL,HAM,HAMAB,HAO,HAS,HAST,HASU, HAW, HB, HBG, HBR, HCIB, HEID, HGI, HIP, HKU, HNHM, HNT, HO, HPL, HRCB, HRP,HSS,HSU,HU,HUA,HUAA,HUAL,HUAZ,HUEFS,HUEM,HUFU,HUSA,HUT, HXBH, HYO, IA, IAA, IAC, IAL, IAN, IB, IBGE, IBUG, ICEL, ICN, IEB, IFO, ILL, ILLS, IMSSM, INB, INEGI, INIF, INM, INPA, IPA, IPRN, ISC, ISL, ISTC, ISU, ITCV, ITMH, IZAC, IZTA, JACA, JBAG, JE, JEPS, JOE, JUA, JYV, K, KANU, KIEL, KMN, KMNH, KOELN,KOR,KPM,KSC,KSTC,KSU,KTU,KU,KUO,KYO,L,LA,LAE,LAF,LAM, LCR, LD, LE, LEB, LEMA, LG, LI, LIL, LINN, LISE, LISI, LISU, LKHD, LL, LM, LOJA, LOMA, LP, LPAG, LPB, LPD, LPS, LSU, LTR, LY, LYJB, LZ, M, MA, MAF, MAIC, MAK, MAN, MARY, MASS, MB, MBK, MBM, MBML, MCM, MCN, MCNS, MEL, MEN, MERL, MEXU, MFA, MFU, MG, MGC, MICH, MIL, MIN, MISS, MJG, MMMN, MNHM, MNHN, MO, MOL, MOR, MOSS, MPU, MPUC, MRSN, MSB, MSC, MSE, MSTR, MSUN, MT, MTMG, MU, MUB, MUCV, MVFA, MVFQ, MVJB, MVM, MY, N, NA, NCSC, NCU, ND, NE, NEB, NHM, NHMC, NHT, NLH, NLU, NMB, NMC, NMCR, NMNL, NMR, NMSU, NMW, NO, NOU, NRCC, NSPM, NSW, NT, NUM, NWOSU, NY, 0, OC, OCLA, ODU, OHN, OKL, OKLA, OMA, OS, OSA, OSC, OSH, OSN, OULU, OWU, OXF, P, PACA, PAR, PE, PEL, PENN, PERTH, PEUFR, PFC, PH, PI, PKDC, PLAT, PMA, PMNH, PNH, POLL, POM, PORT, PR, PRC, PRE, PTBG, PVNH, PY, QCA, QCNE, QFA, QM, QMEX, QRS,QUE,R,RAS,RB,RBR,REG,RENO,RFA,RIOC,RM,RNG,ROST,RPM,RSA, RYU, S, SALA, SAM, SAN, SANT, SAPS, SASK, SBT, SD, SEL, SEV, SF, SFSU, SGO, SI, SIM, SING, SIU, SJRP, SLPM, SMB, SMDB, SMF, SNM, SOM, SP, SPF, SPSF, SQF, SRFA, STL, STU, SUVA, SVG, SZU, TAES, TAI, TAIF, TAMU, TAN, TEF, TENN, TEPB, TEX, TFC, TFM, TI, TKPM, TNS, TO, TRA, TRH, TROM, TRT, TRTC, TRTE, TRTS, TS, TSM, TTRS, TU, TULS, TUR, U, UADY, UAM, UAMIZ, UARK, UAS, UAT, UB, UBA, UBC, UC, UCAM, UCBG, UCR, UEC, UESC, UFG, UFMA, UFMT, UFP, UFRJ, UFRN, UFS, UGDA, UH, UI, UJAT, ULM, ULS, UME, UMO, UNA, UNB, UNCC, UNEX, UNL, UNM, UNR, UNSL, UPCB, UPEI, UPNA, UPNG, UPS, US, USAS, USJ, USM, USNC, USON, USP, USZ, UT, UTC, UTEP, UTMC, UV, UVIC, UVSC, UWO, V, VA, VAL, VALD, VDB, VEN, VM, VMSL, VT, W, WAG, WAT, WELT, WFU, WII, WIN, WIS, WMNH, WOH, WRSL, WS, WTU, WU, XAL, Y, YA, YAM, YU, Z, ZMT, ZSS, ZT.

## Author contributions

TC and SH designed the study, developed the methods and wrote the manuscript. TC compiled the data, performed the analyses and led the writing. UE, HK, AS and PW contributed data and helped refine the study design. AS, HK, PW, SH, UE and HK reviewed and edited the manuscript.

## Data availability statement

Data sharing not applicable to this article as no new datasets were generated during the current study.

## Code availability statement

Computer code to execute the TTR-SDM has been uploaded for the reviewers. We will make the code public on github.com upon acceptance of the manuscript.

## Extended data figure and tables

**Figure El.**
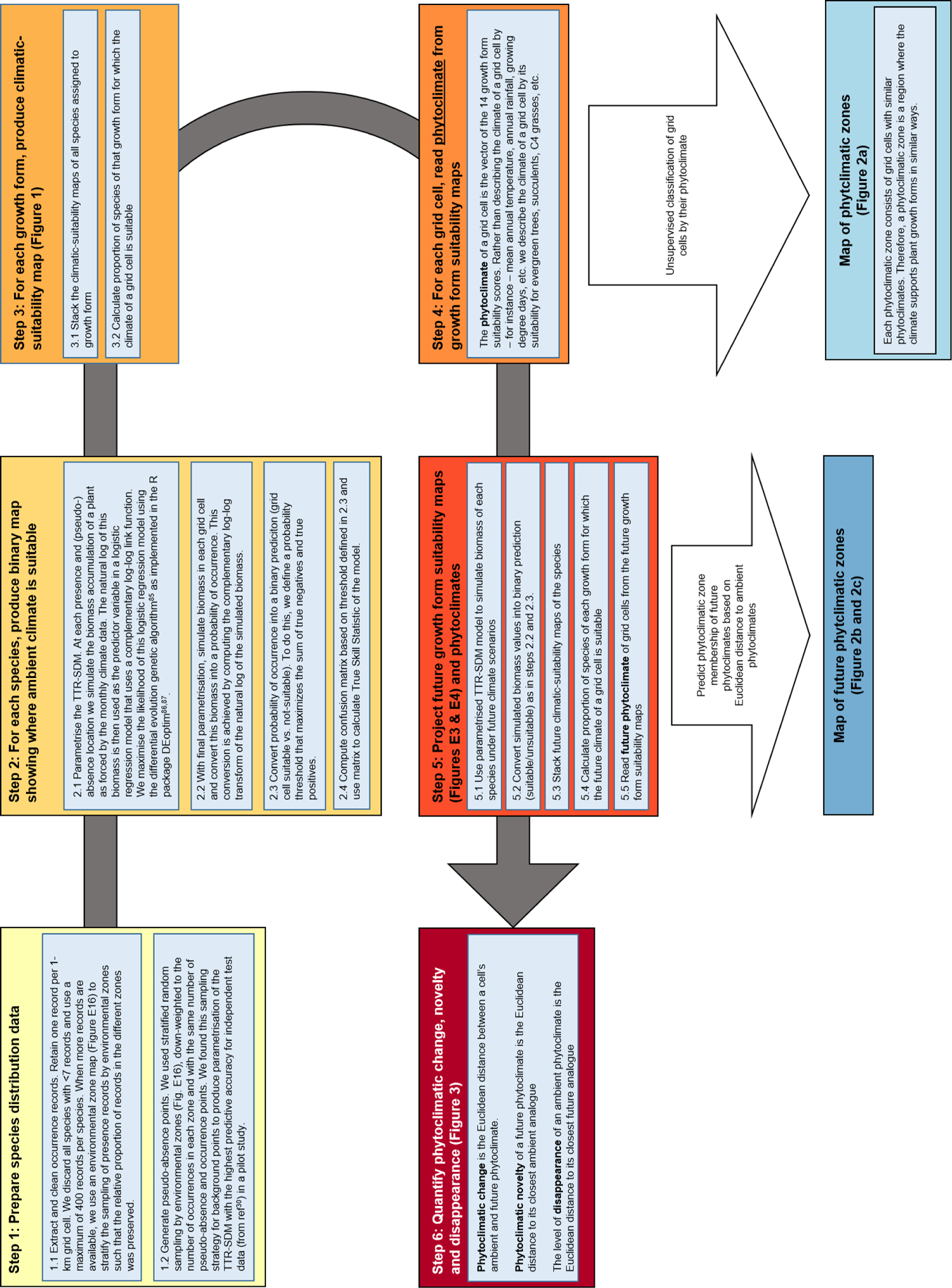
Workflow for analyzing phytoclimates and their change. The original protocol is described in ref^24^.

**Figure E2.**
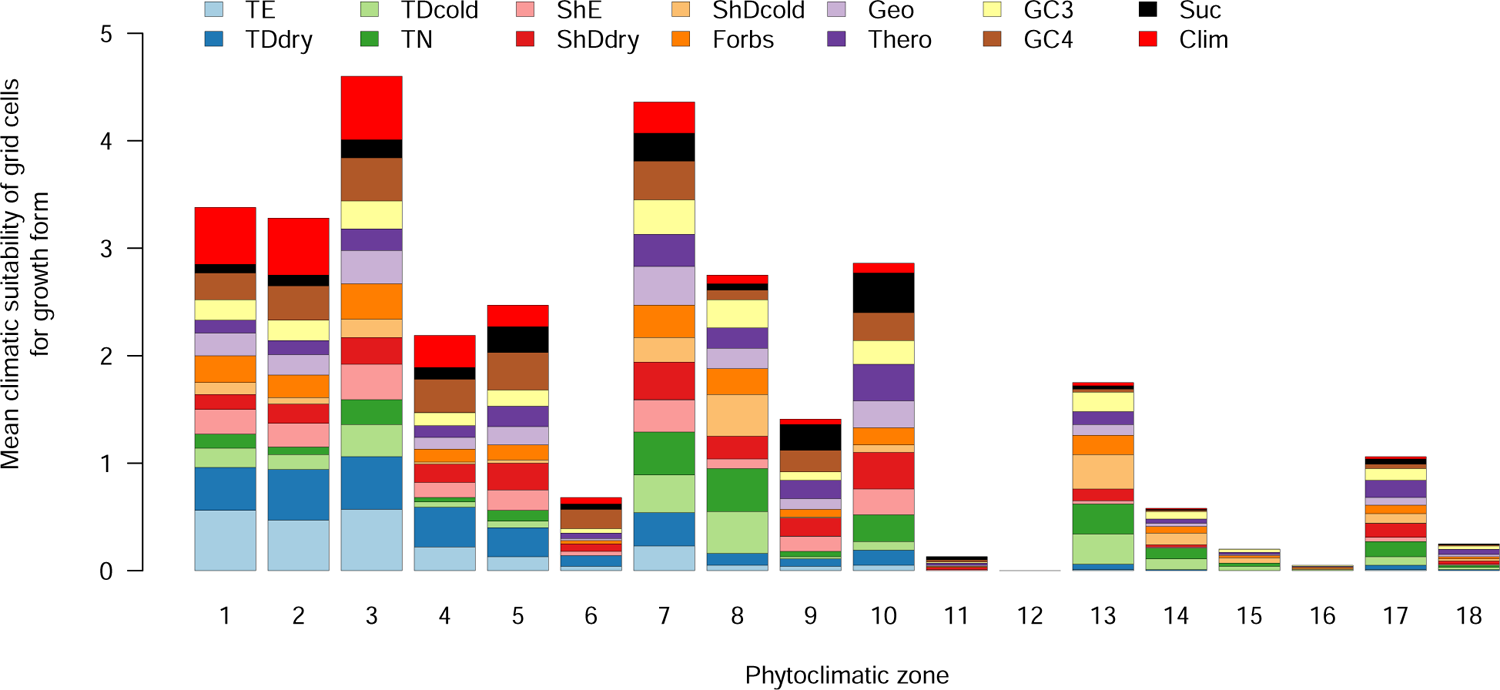
Mean climatic suitability of grid cells for plant growth forms in phytoclimatic zones. The zone numbers are the same as in Fig. 2. The values used to produce the figure are taken from Table E1. TE=evergreen trees, TDdry=drought-deciduous trees, TDcold=cold-deciduous trees, TN=needleleaf trees, ShE=evergreen shrubs, ShDdry=drought-deciduous shrubs, ShDcold=cold-deciduous shrubs, Geo=geophytes, Thero=therophytes, GC3=C3 grasses, GC4=C4 grasses, Suc=succulents, Clim=climbers.

**Figure E3.**
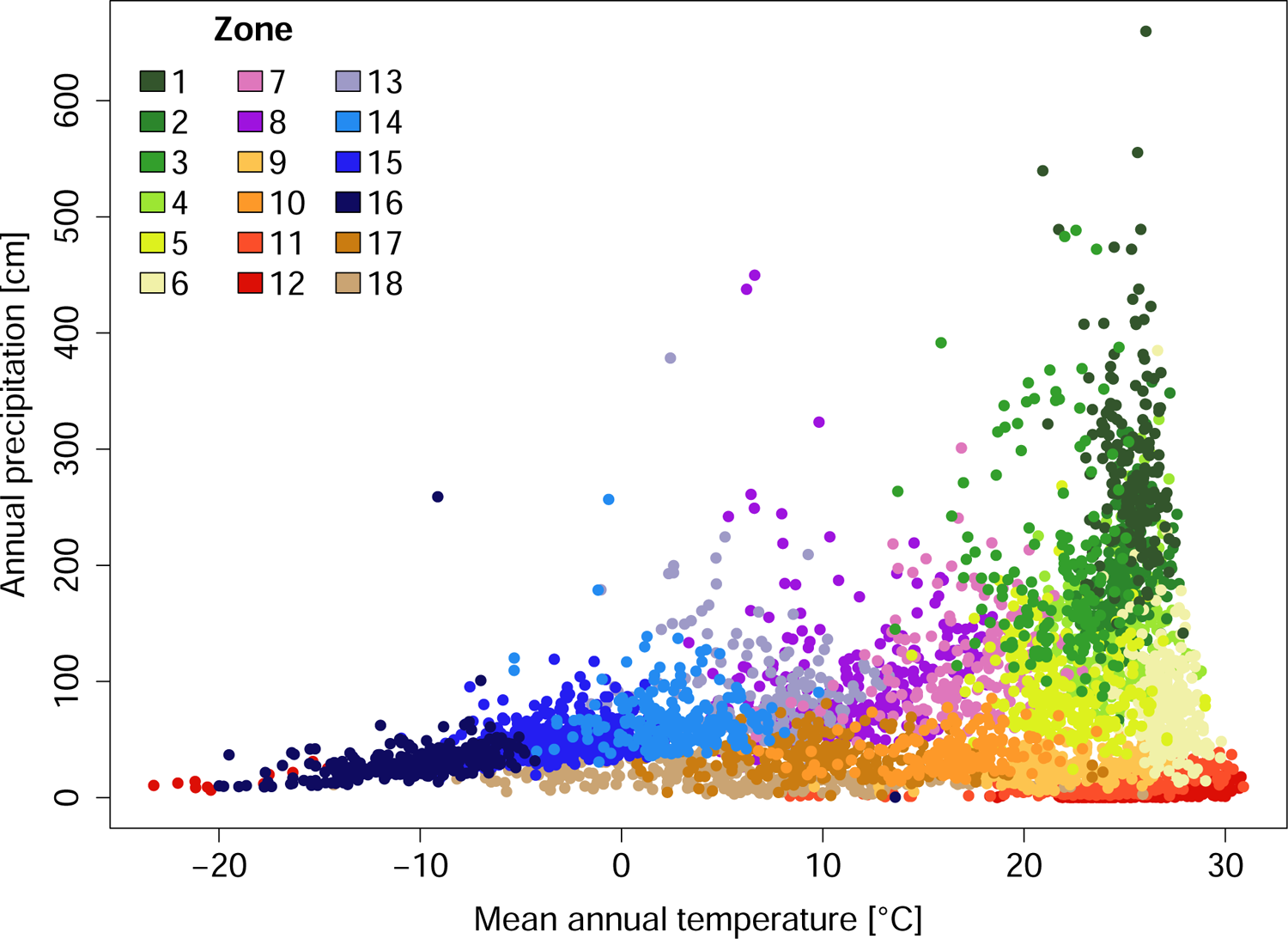
Mean annual temperature and annual precipitation in the phytoclimatic zones. The figure uses a sample of 2.5% of the cells of each phytoclimatic zone. Zone 12 includes grid cells in the high arctic tundra and extreme dry and hot deserts, both of which have extremely low climatic suitability for all growth forms. Therefore, these cells are in the same phytoclimatic zone.

**Figure E4.**
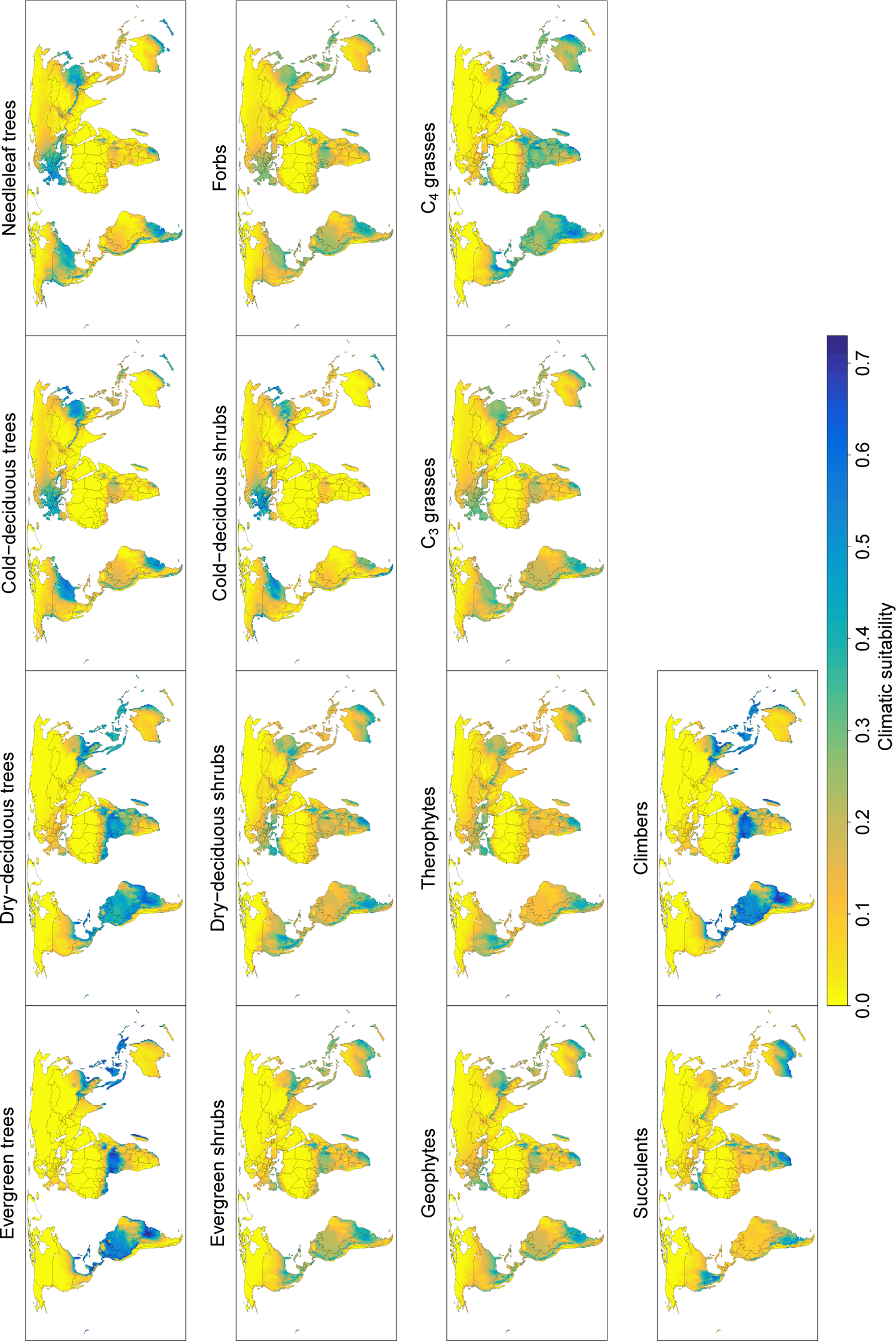
Climatic suitability for major plant growth forms in 2070 under RCP 2.6. Suitability is expressed as the proportion of plant species of a growth form for which the cells’ climate is suitable according to an ecophysiological plant growth model. The values are means across climatologies projected by five Global Circulation Models. Note that the range of the legend is slightly larger than that in Fig. 1 in the main text.

**Figure E5.**
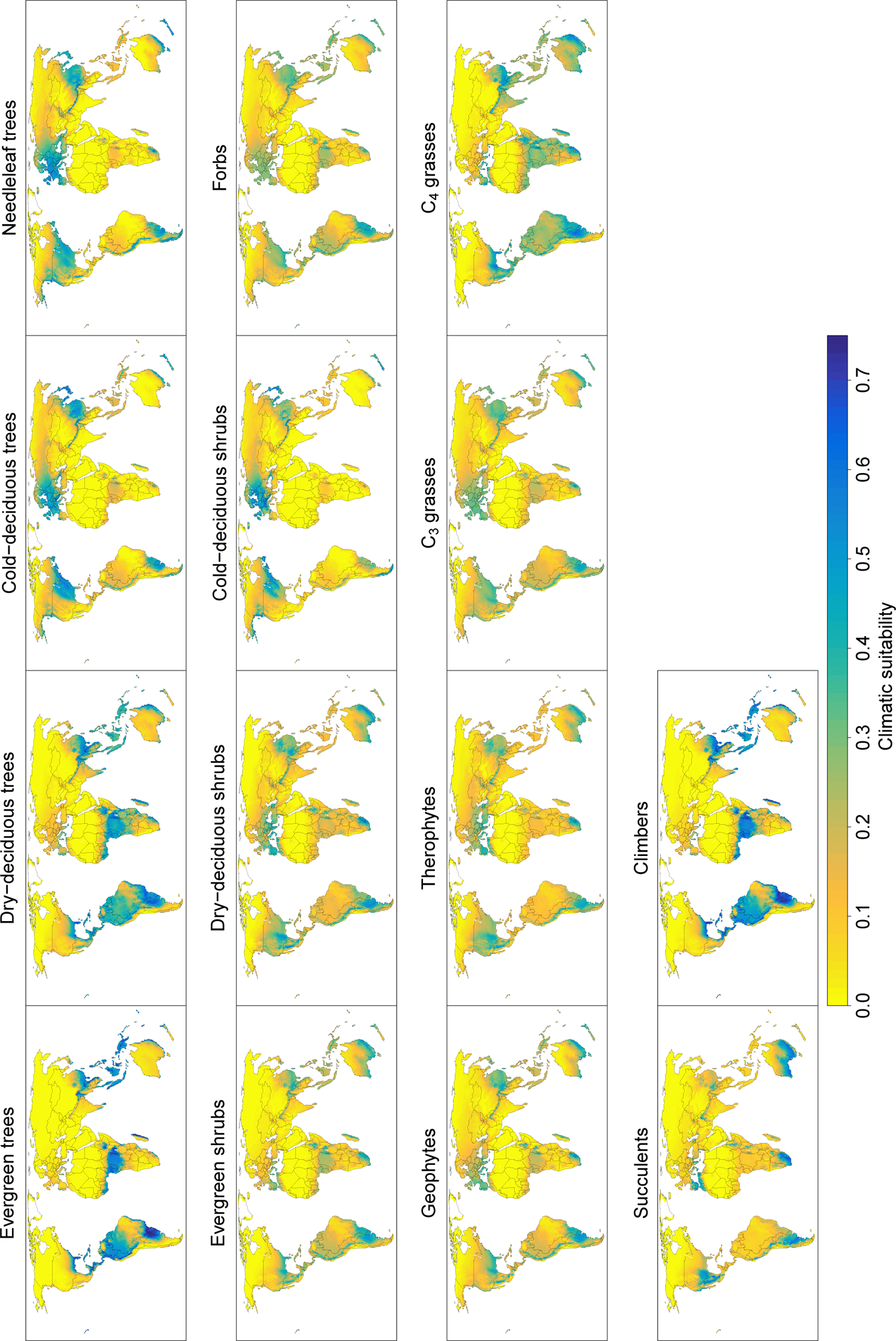
Climatic suitability for major plant growth forms in 2070 under RCP 8.5. Suitability is expressed as the proportion of plant species of a growth form for which the cells’ climate is suitable according to an ecophysiological plant growth model. The values are means across climatologies projected by five Global Circulation Models. Note that the range of the legend is slightly larger than that in Fig. 1 in the main text.

**Figure E6.**
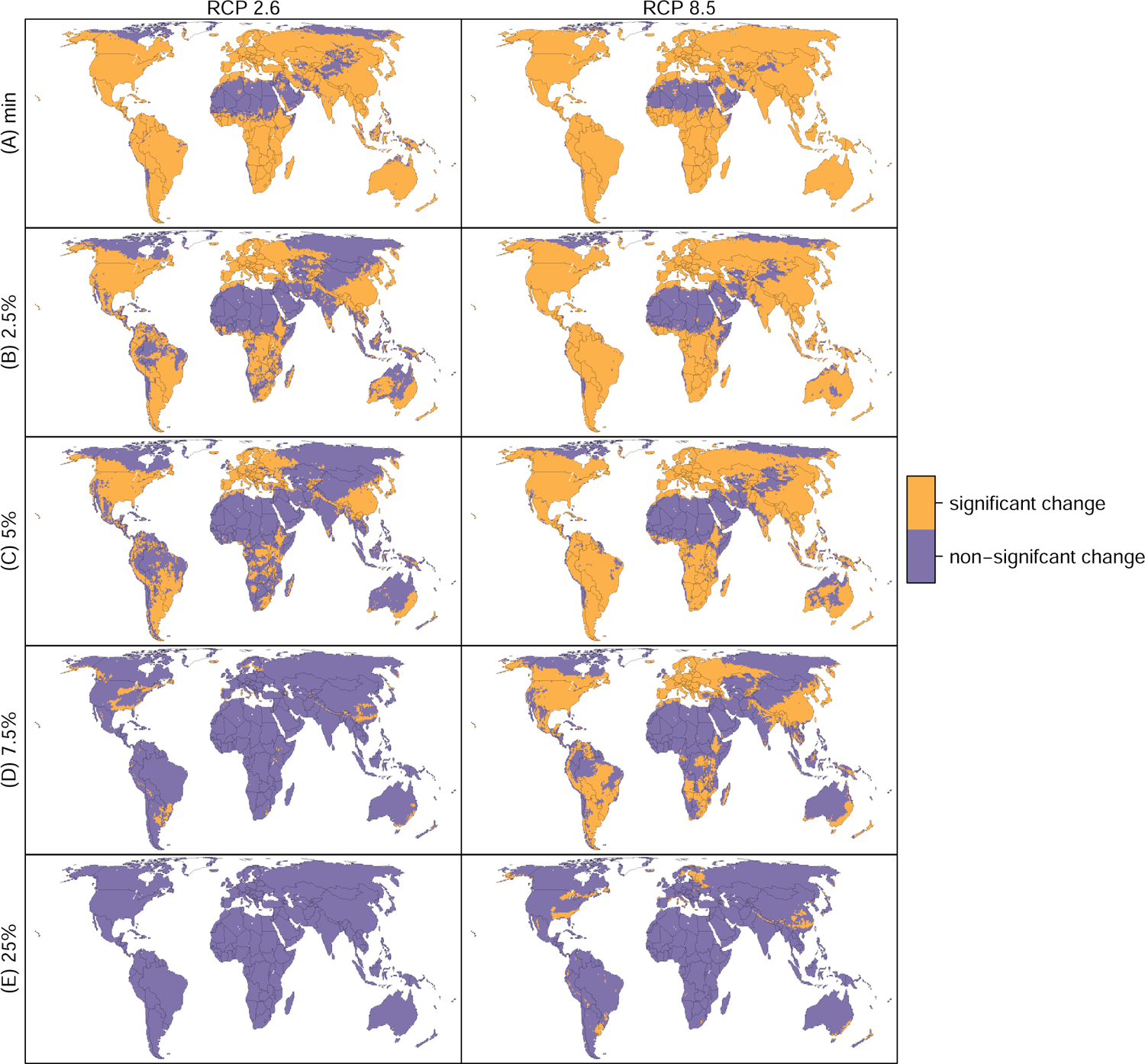
Regions with significant phytoclimatic change when different threshold values for assessing significance are applied. The threshold values are (A) the distance between the centroids of the two most similar phytoclimatic zones in Euclidean growth form suitability space, (B) the 2.5^th^ percentile, (C) the 5^th^ percentile, (D) the 7.5^th^ percentile, and (E) the 25^th^ percentile of the pairwise inter-centroid distances between the phytolcimatic zones.

**Figure E7.**
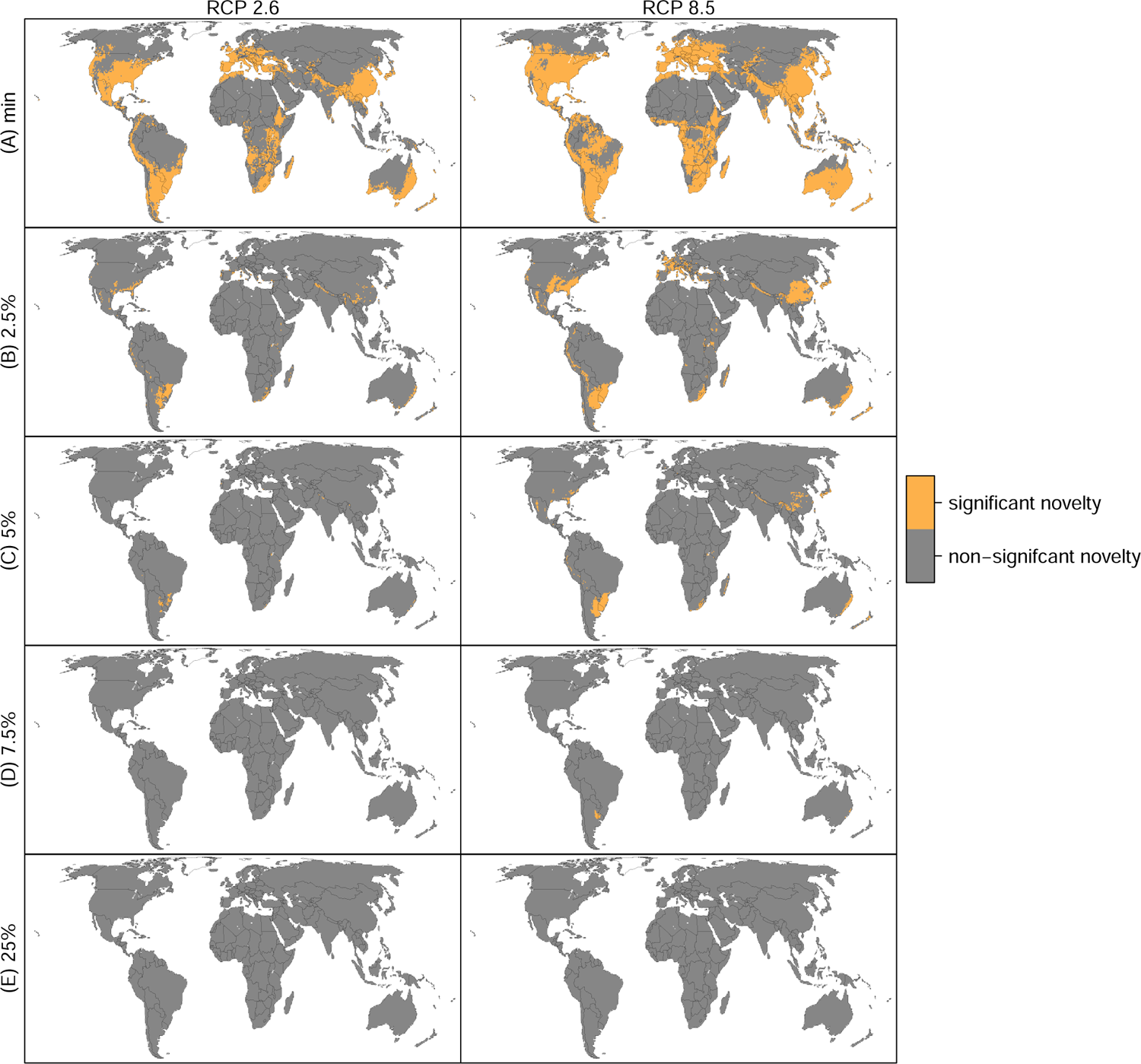
Regions with significant phytoclimatic novelty when different threshold values for assessing significance are applied. The threshold values are (A) the distance between the centroids of the two most similar phytoclimatic zones in Euclidean growth form suitability space, (B) the 2.5^th^ percentile, (C) the 5^th^ percentile, (D) the 7.5^th^ percentile, and (E) the 25^th^ percentile of the pairwise inter-centroid distances between the phytolcimatic zones.

**Figure E8.**
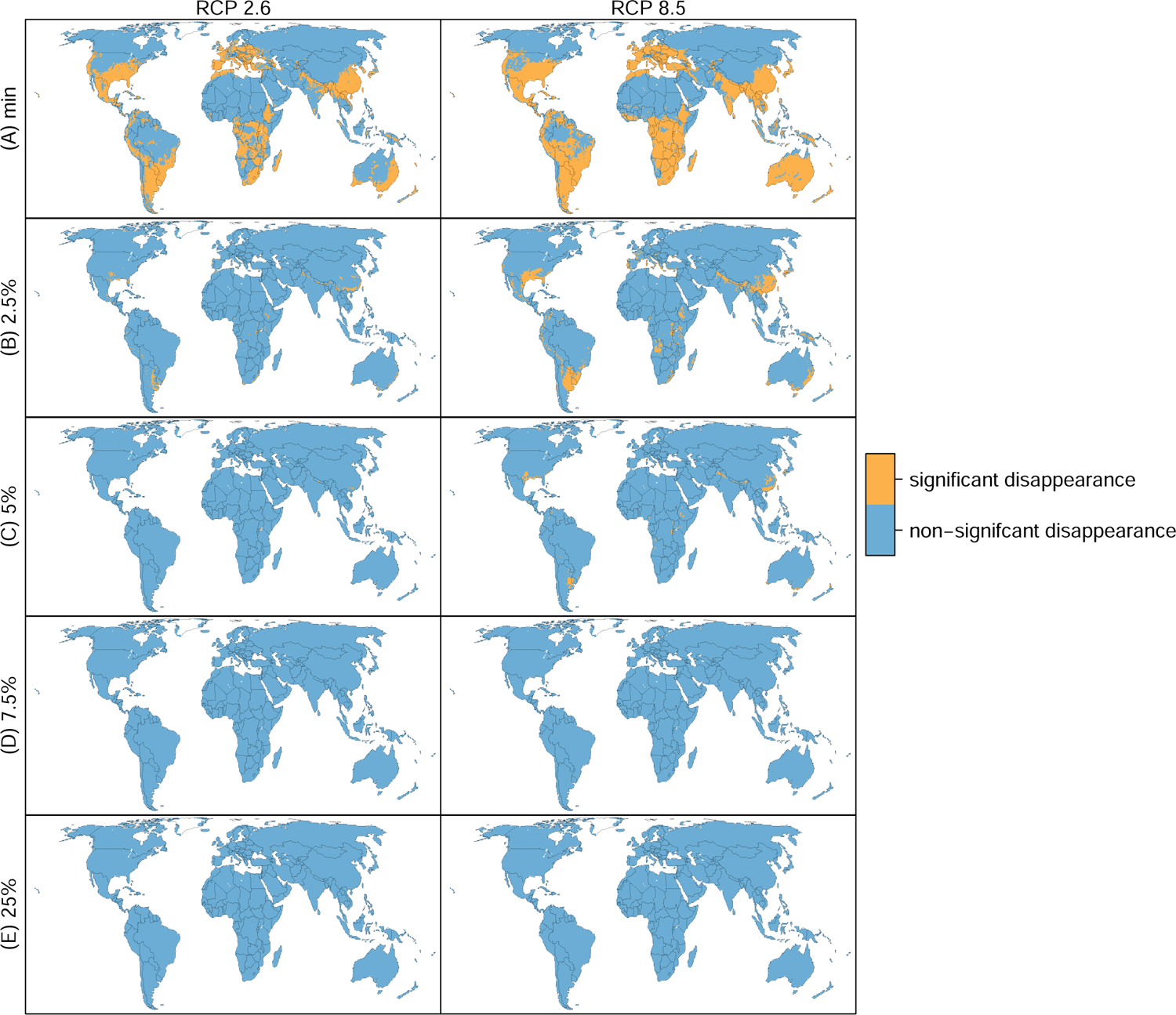
Regions with significant disappearance of phytoclimate when different threshold values for assessing significance are applied. The threshold values are (A) the distance between the centroids of the two most similar phytoclimatic zones in Euclidean growth form suitability space, (B) the 2.5^th^ percentile, (C) the 5^th^ percentile, (D) the 7.5^th^percentile, and (E) the 25^th^ percentile of the pairwise inter-centroid distances between the phytolcimatic zones.

**Figure E9.**
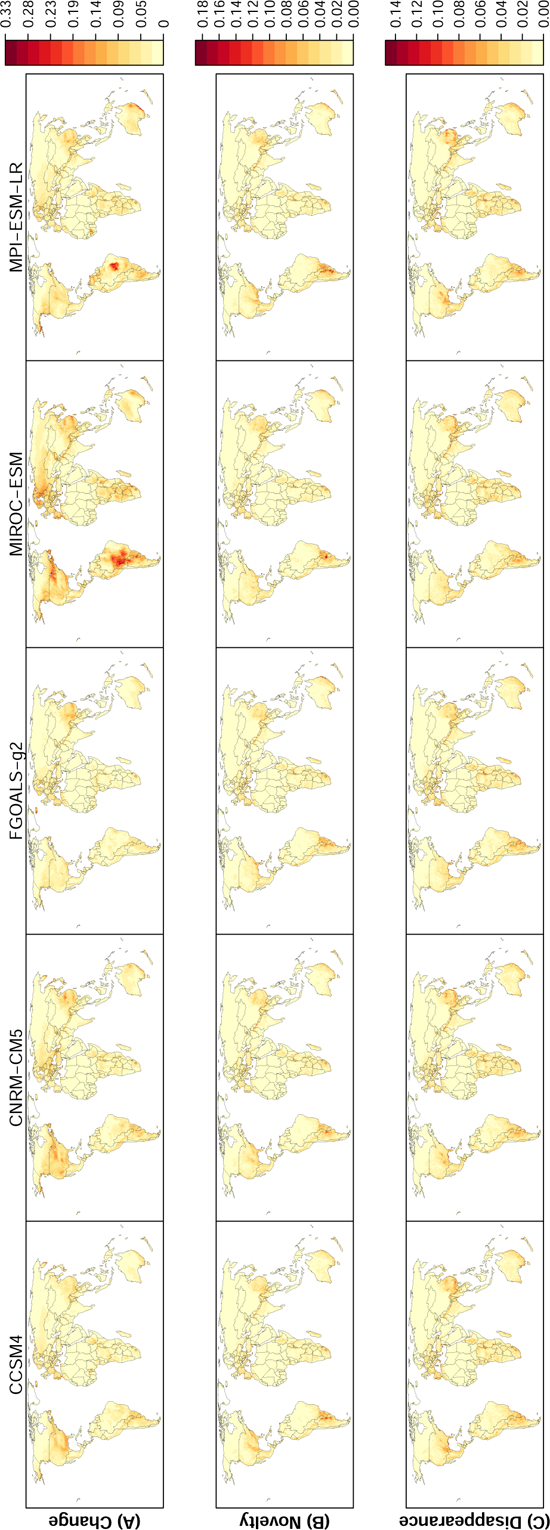
(A) Change, (B) novelty and (C) disappearance of phytoclimates by 2070 under RCP 2.6 for five different Global Circulation Models. The phytoclimate of the cells is the suitability of the local climate for 14 plant growth forms that characterise the structure of terrestrial ecosystems. (A) Local change in phytoclimate. (B) Risk of novelty of the projected phytoclimate in 2070, expressed as the Euclidean distance of a cell’s future phytoclimate to its closest ambient analogue. (C) Risk of disappearance of the existing phytoclimate, expressed as the Euclidean distance of a cell’s ambient phytoclimate to its closest future analogue.

**Figure E10.**
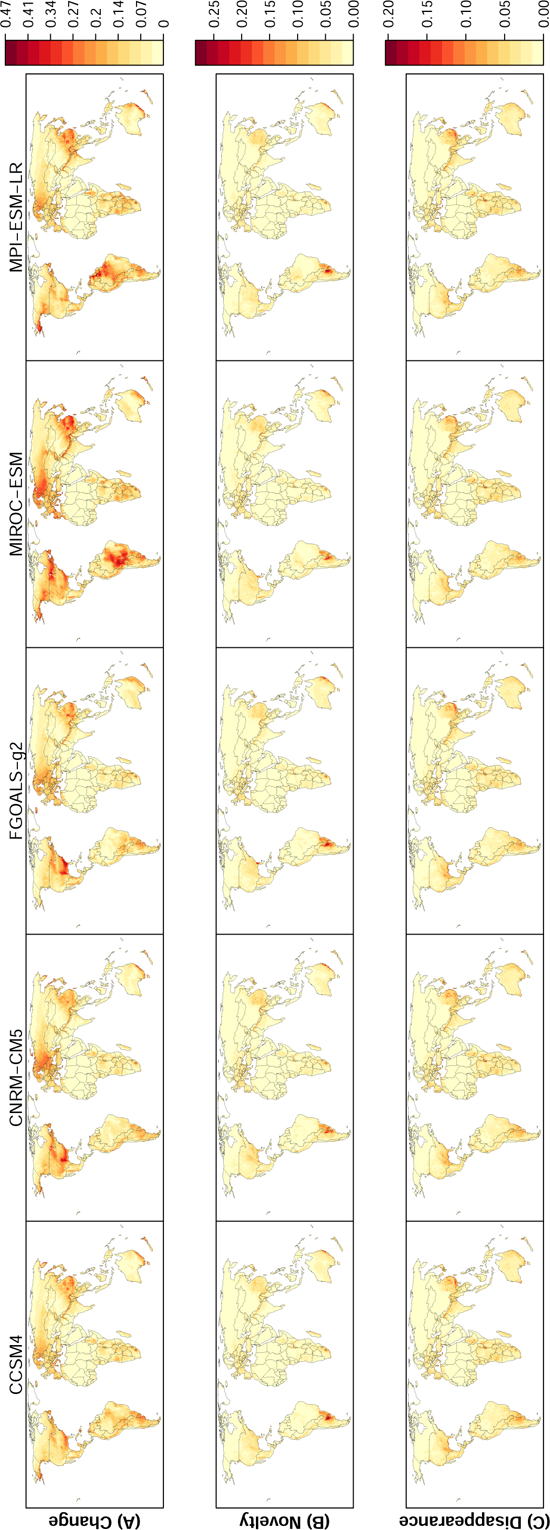
(A) Change, (B) novelty and (C) disappearance of phytoclimates by 2070 under RCP 8.5 for five different Global Circulation Models. The phytoclimate of the cells is the suitability of the local climate for 14 plant growth forms that characterise the structure of terrestrial ecosystems. (A) Local change in phytoclimate. (B) Risk of novelty of the projected phytoclimate in 2070, expressed as the Euclidean distance of a cell’s future phytoclimate to its closest ambient analogue. (C) Risk of disappearance of the existing phytoclimate, expressed as the Euclidean distance of a cell’s ambient phytoclimate to its closest future analogue.

**Figure E11.**
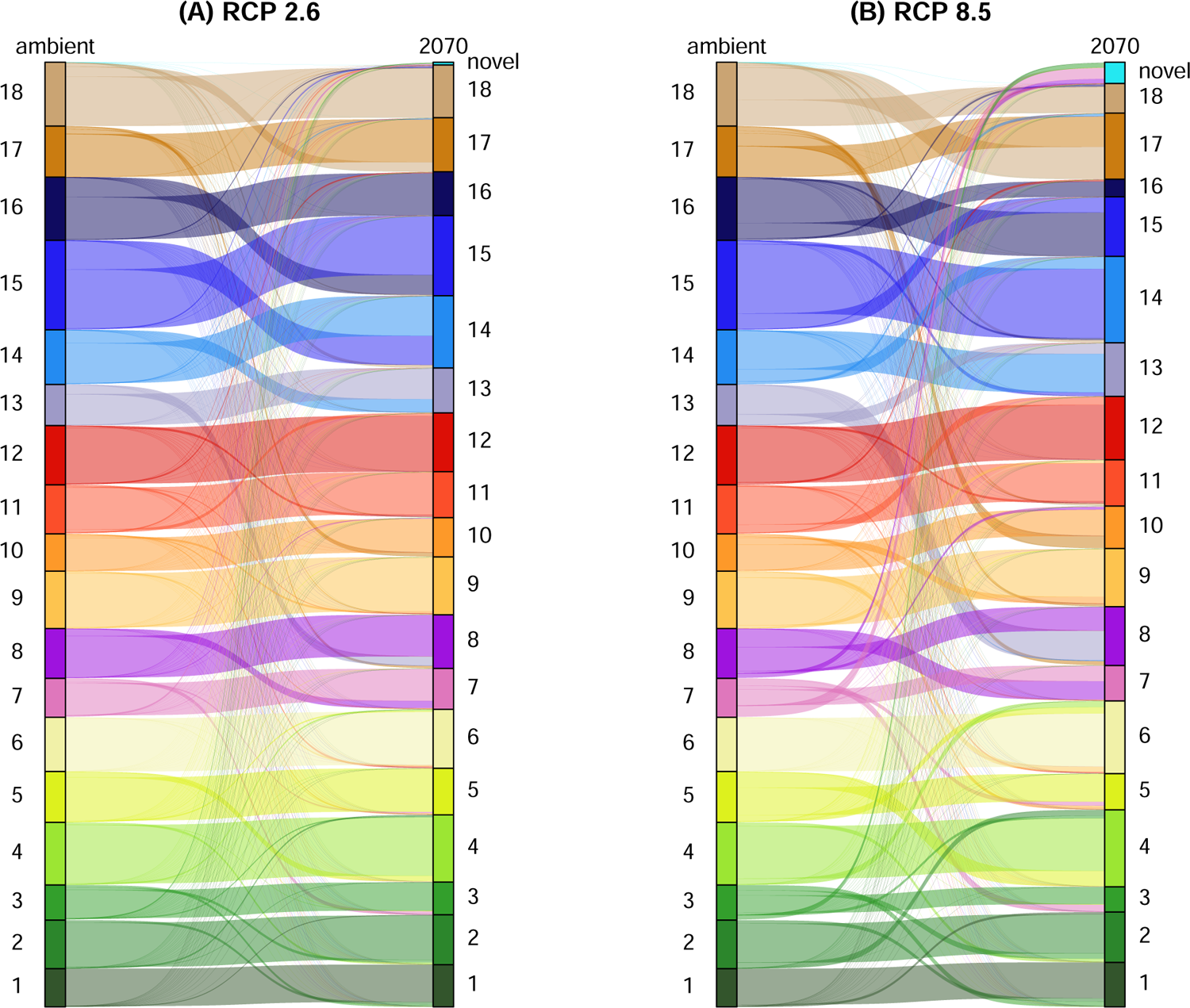
Projected phytoclimatic zone transitions of grid cells by 2070 under (A) RCP 2.6 and (B) RCP 8.5. Zone numbers are the same as in Fig. 2. The median growth form suitabilities of each grid cell in 2070 across projections from five Global Circulation were used to asses the future zone membership of each cell.

**Figure El2.**
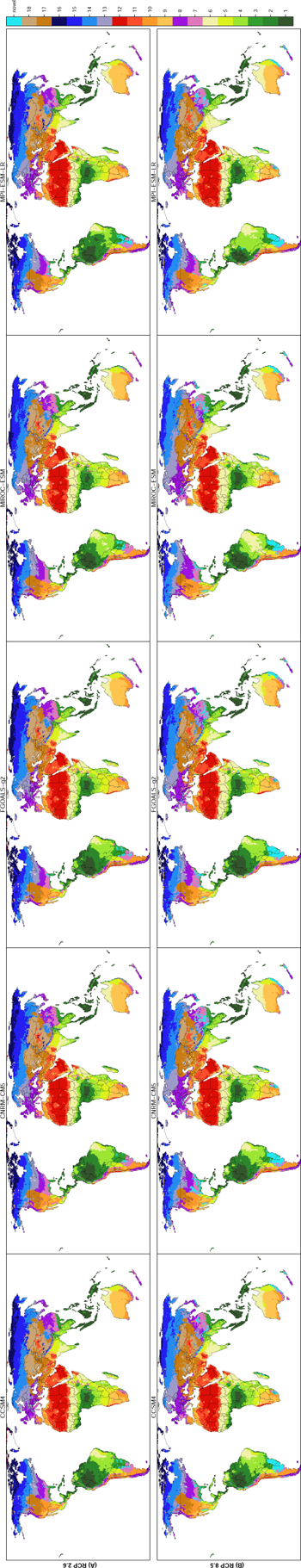
Projected distribution of phytoclimatic zones in 2070 under (A) RCP 2.6 and (B) RCP 8.5 when using climate data from five different Global Circulation Models.

**Figure E13.**
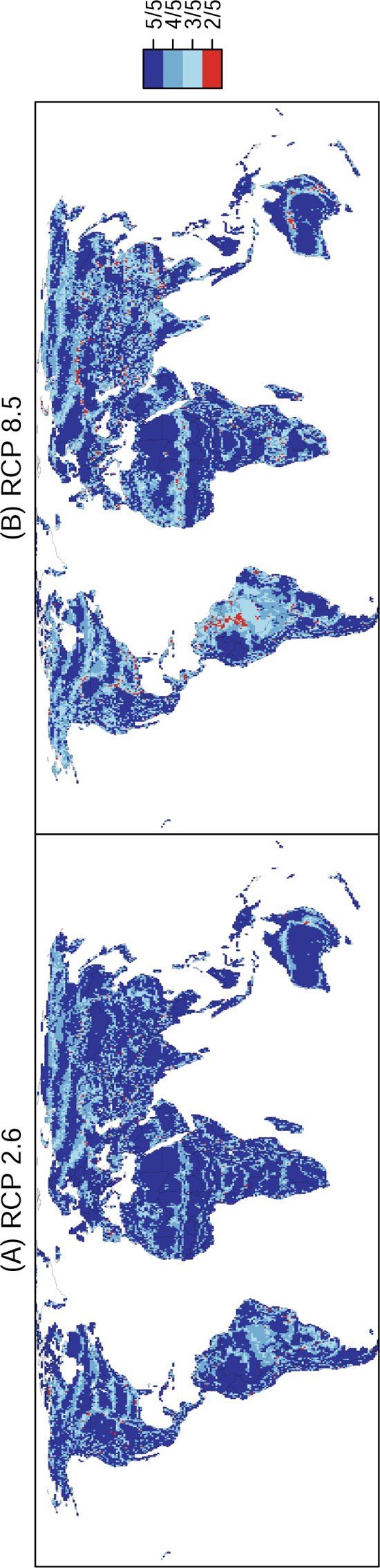
Agreement on phytoclimatic zone in 2070 between five projections that used 2070 climate data from different Global Circulation Models. A value of 5/5 means that all five projections made the same 2070 phytoclimatic zone prediction. A value of 2/5 means that only two of five projections made the same 2070 phytoclimatic zone prediction.

**Figure E14.**
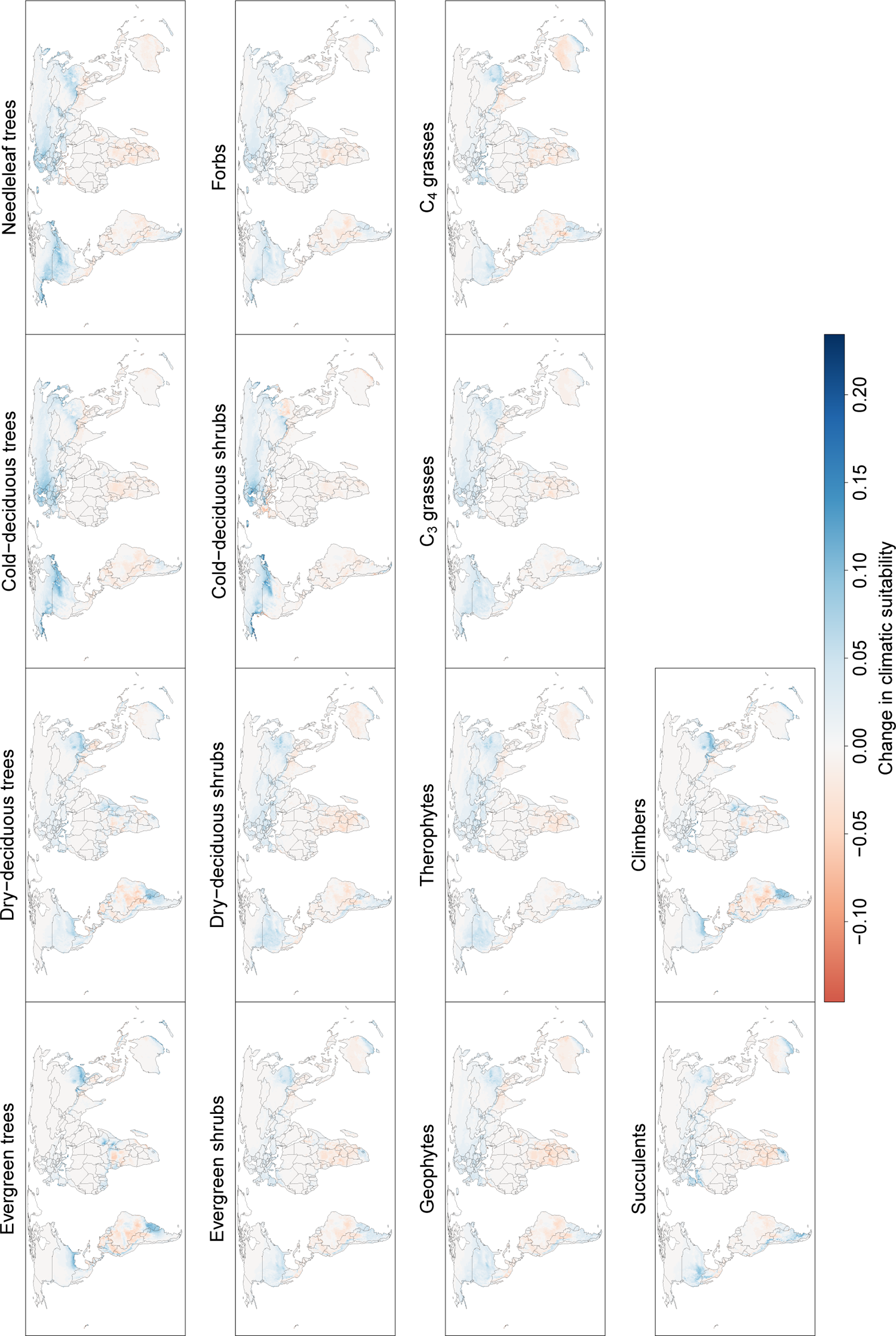
Change in climatic suitability of grid cells for plant growth forms between ambient and 2070 climatologies under RCP 2.6. Change is expressed as change in proportion of species of a growth form that can grow in a cell according to our simulations. Values are medians across five projections that used 2070 climate data from different Global Circulation Models.

**Figure E15.**
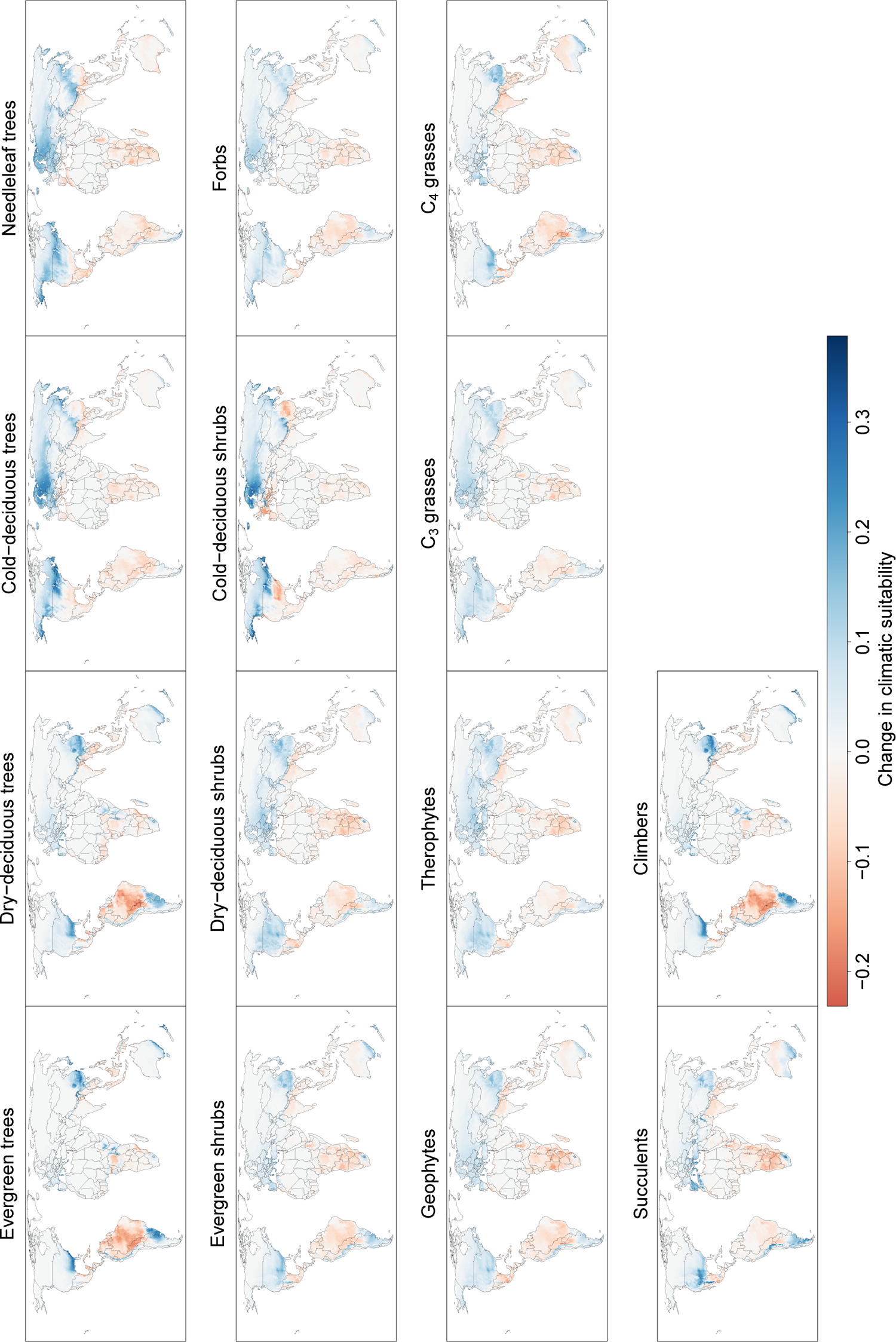
Change in climatic suitability of grid cells for plant growth forms between ambient and 2070 climatologies under RCP 8.5. Change is expressed as change in proportion of species of a growth form that can grow in a cell according to our simulations. Values are medians across five projections that used 2070 climate data from different Global Circulation Models.

**Figure E16.**
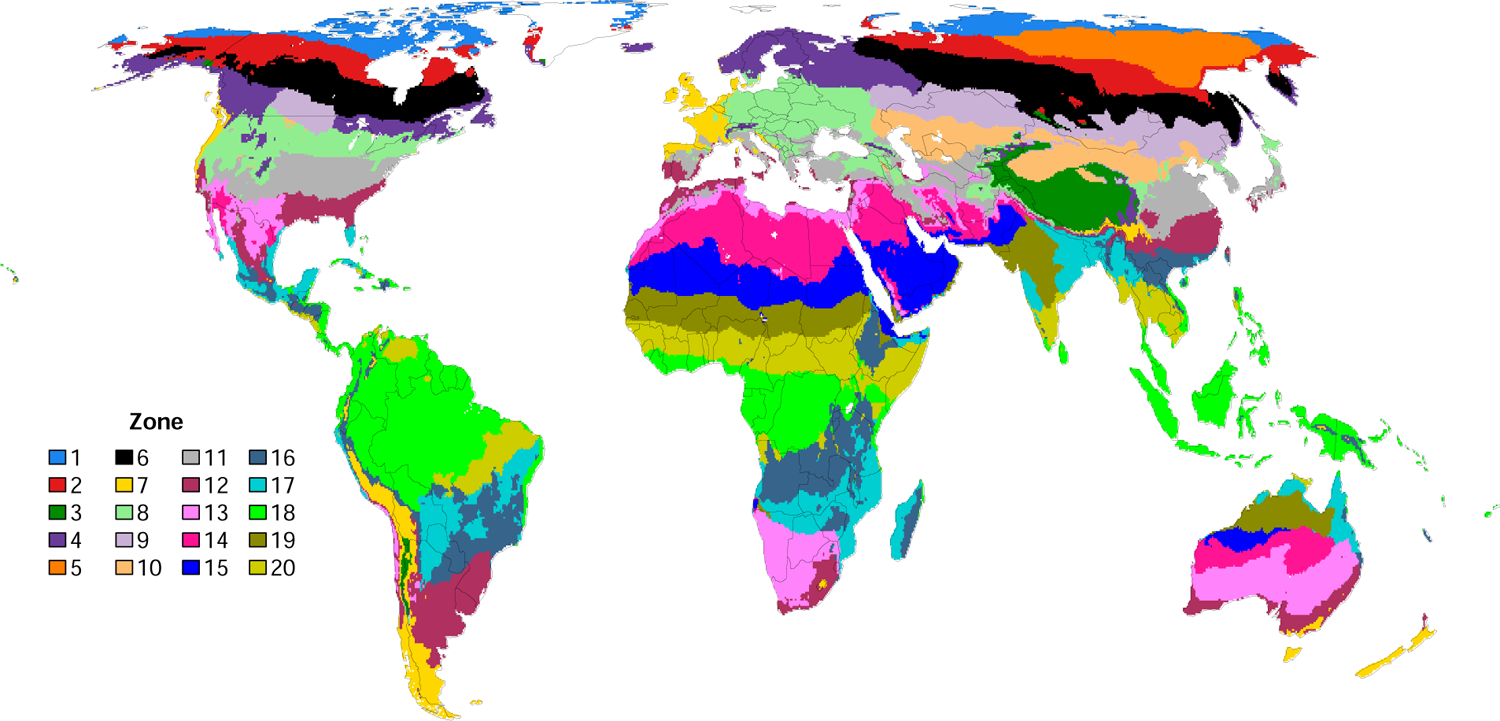
Environmental zones used to stratify the sampling of presence and pseudo-absence points for species distribution modelling. The zones were obtained from a classification of grid cells based on the input variables of the TTR,-SDM, i.e. monthly values of mean., maximum and minimum temperature., soil water content and solar radiation. The TTR,..SDM also uses atmospheric CO2 concentration, but CO2 was assumed to be equally distributed globally and was thus not used in this classification of grid cells.

**Figure E17.**
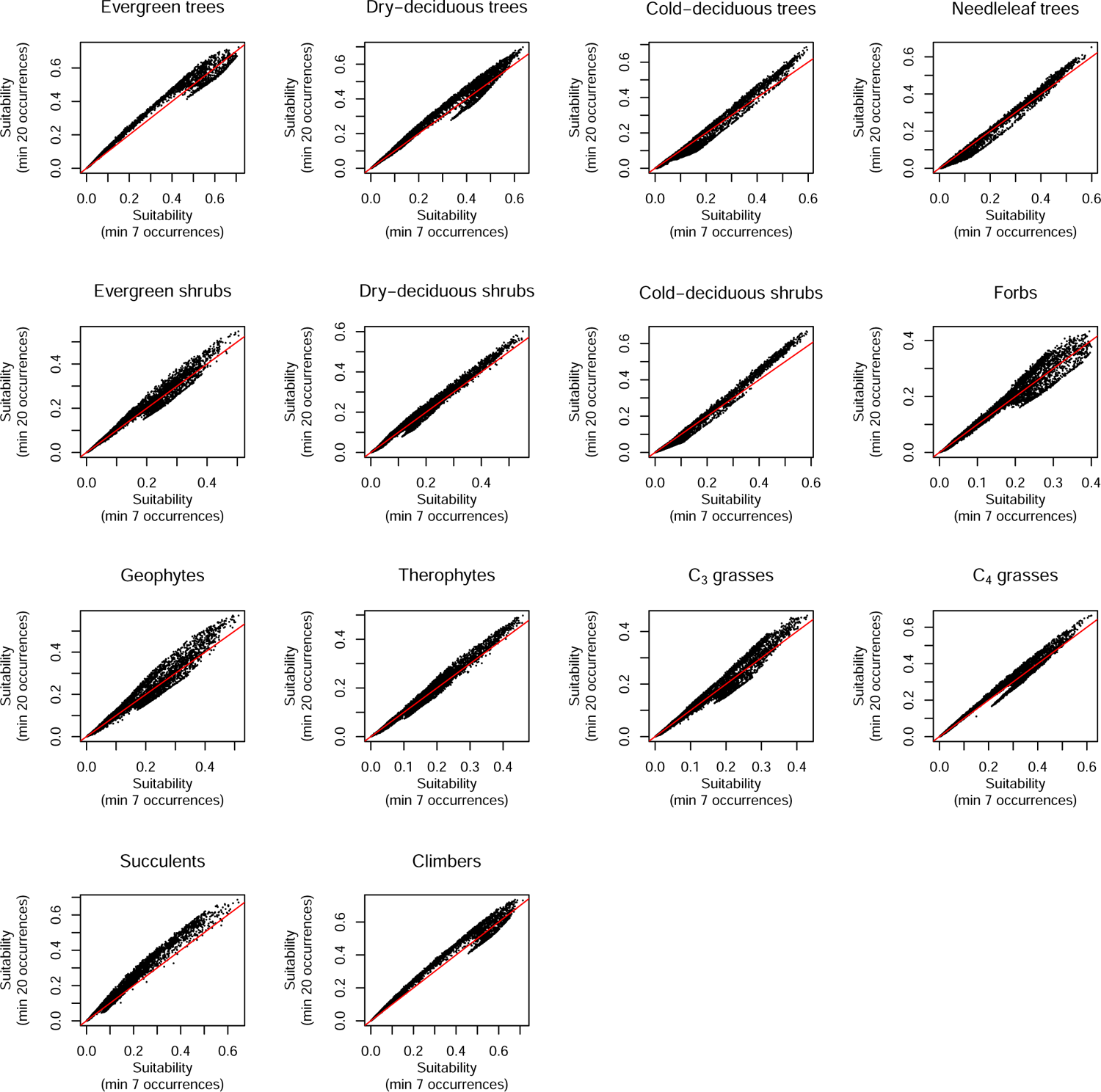
Modelled climatic suitability of grid cells for plant growth forms derived from range models with a **minimum** of seven versus 20 occurrence records.

**Figure E18.**
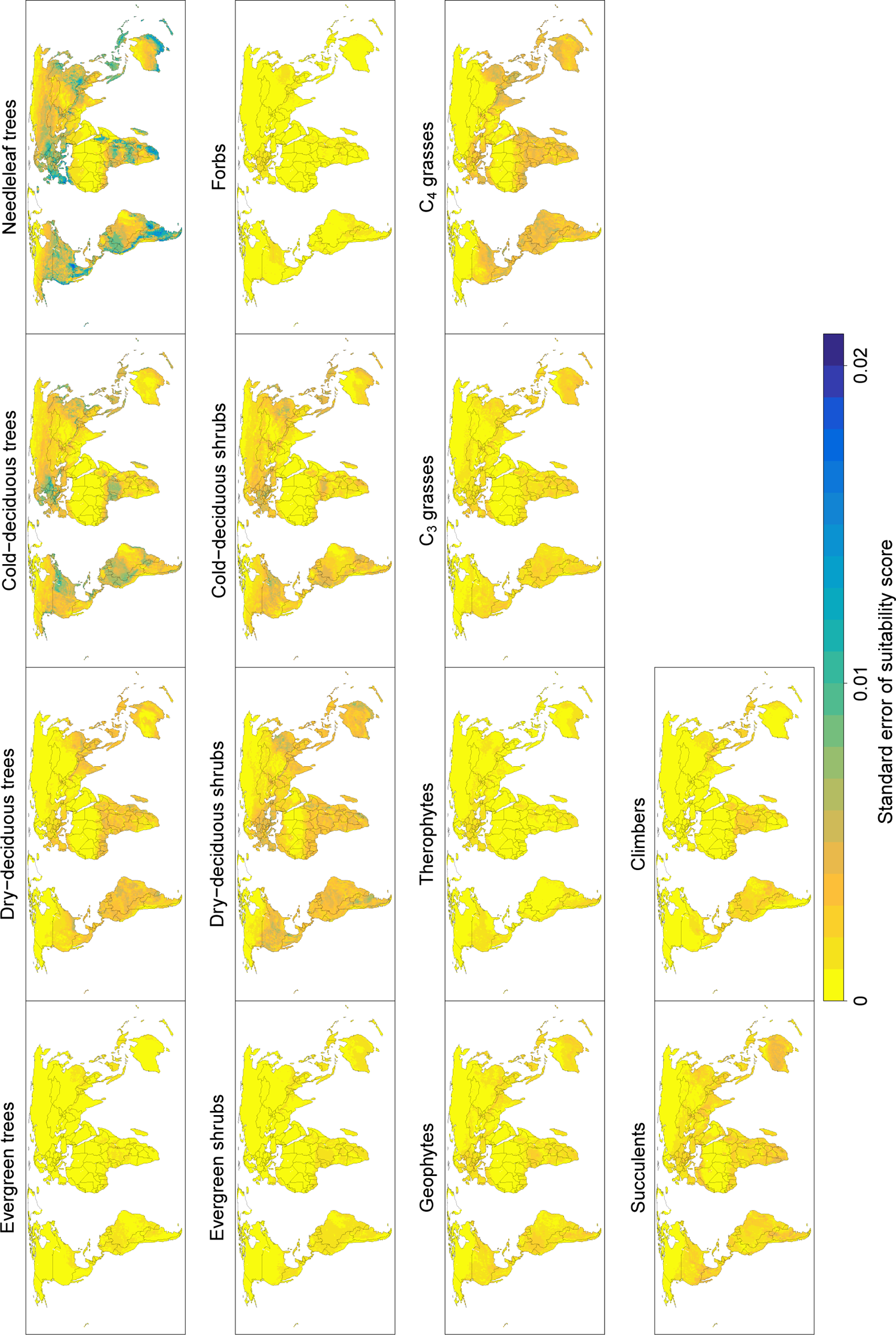
Standard error of suitability values calculated from five random subsets of 50% of the species from each growth form. The suitability values can range between 0 and 1. The values of the standard errors therefore indicate that the suitability estimate are accurate. The highest standard errors occur in the needleleaf trees, the total number of which was 439.

**Figure E19.**
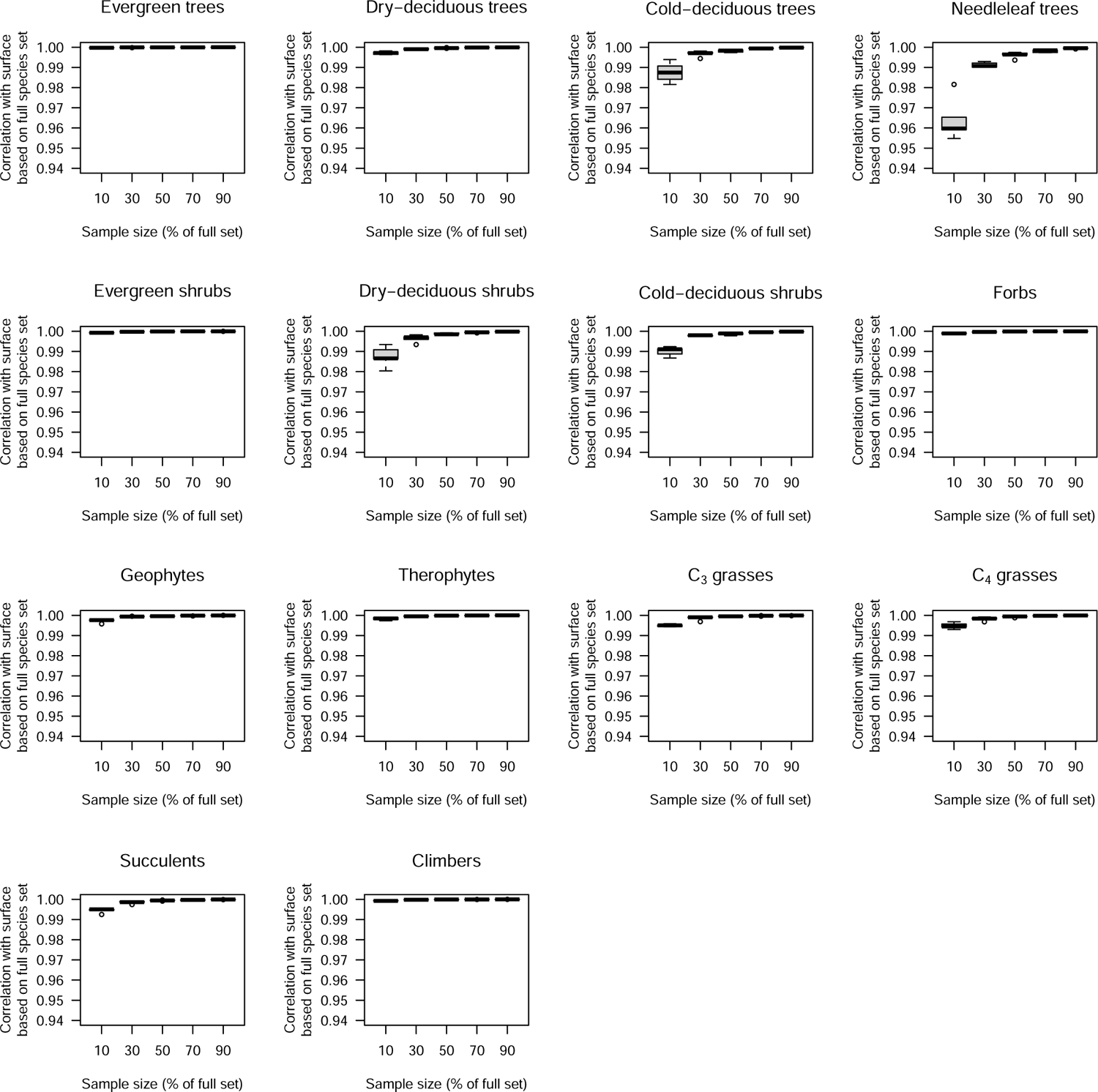
Correlations of growth form suitability values estimated based on random subsets of species with the growth form suitability values based on the full set of species. Boxplots show spread of correlations across five replicate random subsets. The correlations with the full-set suitabilities approximate 1 already when smaller subsets are used, indicating that adding further species would not change the suitability estimates any further.

**Figure E20.**
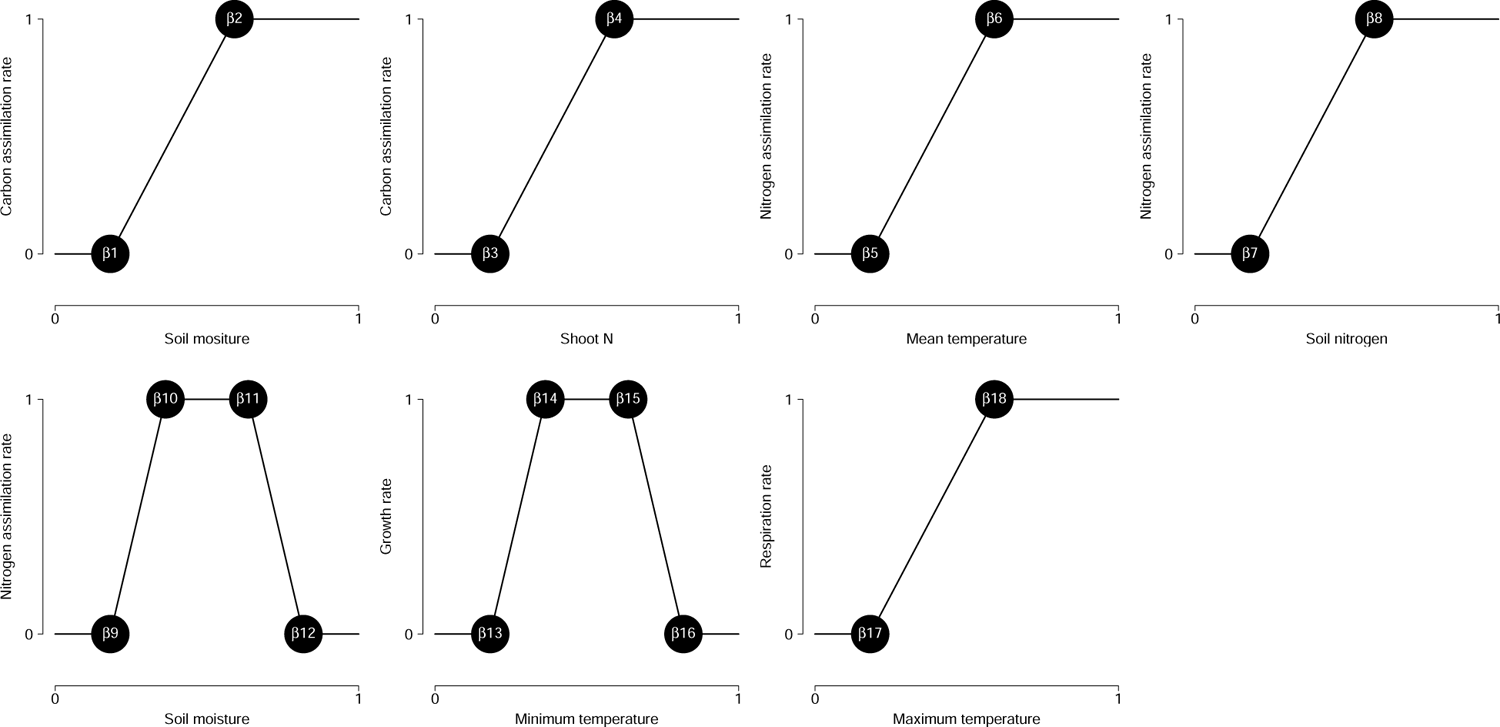
Graphical representation of the parameters of TTR-SDM. The panels show the model’s representation of how carbon and nitrogen uptake rates as well as growth and respiration rates are influenced by environmental variables. The x-axis coordinates of the 18 black dots are the parameters of the model. They vary among species and we estimate them from species distribution data. In addition, a Farquhar photosynthesis model is used to simulate how solar radiation, temperature and atmospheric CO2 concentrations influence the monthly carbon uptake rates^24^. The Farquhar model used the same photosynthesis parameters for all C3 and C4 plants, respectively, and is therefore not shown. In this application of the TTR-SDM we assumed that soil nitrogen is spatially invariant, which means that nitrogen uptake rates are influenced by temperature and soil moisture only. This is appropriate when the goal is to estimate the climatic niche only. Note that carbon uptake is also influenced by shoot nitrogen (second panel), which is used as a proxy for leaf nitrogen, and that shoot nitrogen is not estimated from species distribution data, but modelled as a function of the plant’s nitrogen pools as described in ref^32^. Also note that the numbering of the parameters differs from ref^32^ due to differences in how carbon uptake is modelled.

---

**Table E1.** Mean climatic suitability of grid cells for plant growth forms in phytoclimatic zones. The zone numbers are the same as in Fig. 2. Standard errors of the means were always≤0.002. TE=evergreen trees, TDdry=drought-deciduous trees, TDcold=cold-deciduous trees, TN=needleleaf trees, ShE=evergreen shrubs, ShDdry=drought-deciduous shrubs, ShDcold=cold-deciduous shrubs, Geo=geophytes, Thero=therophytes, GC3=C3 grasses, GC4=C4 grasses, Suc=succulents, Clim=climbers.

| Zone | TE | TDdry | TDcold | TN | ShE | ShDdry | ShDcold | Forbs | Geo | Thero | GC3 | GC4 | Suc | Clim |
| --- | --- | --- | --- | --- | --- | --- | --- | --- | --- | --- | --- | --- | --- | --- |
| 1 | 0.56 | 0.40 | 0.18 | 0.13 | 0.23 | 0.14 | 0.11 | 0.25 | 0.21 | 0.12 | 0.19 | 0.25 | 0.08 | 0.53 |
| 2 | 0.47 | 0.47 | 0.14 | 0.07 | 0.22 | 0.18 | 0.06 | 0.21 | 0.19 | 0.13 | 0.19 | 0.32 | 0.10 | 0.53 |
| 3 | 0.57 | 0.49 | 0.30 | 0.23 | 0.33 | 0.25 | 0.17 | 0.33 | 0.31 | 0.20 | 0.26 | 0.40 | 0.17 | 0.59 |
| 4 | 0.22 | 0.37 | 0.05 | 0.04 | 0.14 | 0.17 | 0.02 | 0.12 | 0.11 | 0.11 | 0.12 | 0.31 | 0.11 | 0.30 |
| 5 | 0.13 | 0.27 | 0.06 | 0.10 | 0.19 | 0.25 | 0.03 | 0.14 | 0.17 | 0.19 | 0.15 | 0.35 | 0.24 | 0.20 |
| 6 | 0.04 | 0.10 | 0.00 | 0.00 | 0.04 | 0.07 | 0.00 | 0.03 | 0.02 | 0.05 | 0.04 | 0.18 | 0.05 | 0.06 |
| 7 | 0.23 | 0.31 | 0.35 | 0.40 | 0.30 | 0.35 | 0.23 | 0.30 | 0.36 | 0.30 | 0.32 | 0.36 | 0.26 | 0.29 |
| 8 | 0.05 | 0.11 | 0.39 | 0.40 | 0.09 | 0.21 | 0.39 | 0.24 | 0.19 | 0.19 | 0.26 | 0.09 | 0.06 | 0.08 |
| 9 | 0.04 | 0.07 | 0.02 | 0.05 | 0.14 | 0.17 | 0.01 | 0.07 | 0.10 | 0.17 | 0.08 | 0.20 | 0.24 | 0.05 |
| 10 | 0.05 | 0.14 | 0.08 | 0.25 | 0.24 | 0.34 | 0.07 | 0.16 | 0.25 | 0.34 | 0.22 | 0.26 | 0.37 | 0.09 |
| 11 | 0.00 | 0.00 | 0.00 | 0.00 | 0.01 | 0.02 | 0.00 | 0.01 | 0.01 | 0.02 | 0.01 | 0.02 | 0.03 | 0.00 |
| 12 | 0.00 | 0.00 | 0.00 | 0.00 | 0.00 | 0.00 | 0.00 | 0.00 | 0.00 | 0.00 | 0.00 | 0.00 | 0.00 | 0.00 |
| 13 | 0.01 | 0.05 | 0.28 | 0.28 | 0.03 | 0.11 | 0.32 | 0.18 | 0.10 | 0.12 | 0.18 | 0.03 | 0.03 | 0.03 |
| 14 | 0.00 | 0.01 | 0.10 | 0.10 | 0.01 | 0.02 | 0.11 | 0.06 | 0.03 | 0.04 | 0.07 | 0.01 | 0.01 | 0.01 |
| 15 | 0.00 | 0.00 | 0.04 | 0.03 | 0.00 | 0.00 | 0.05 | 0.02 | 0.01 | 0.02 | 0.03 | 0.00 | 0.00 | 0.00 |
| 16 | 0.00 | 0.00 | 0.01 | 0.00 | 0.00 | 0.00 | 0.01 | 0.01 | 0.00 | 0.01 | 0.01 | 0.00 | 0.00 | 0.00 |
| 17 | 0.01 | 0.04 | 0.08 | 0.14 | 0.04 | 0.13 | 0.09 | 0.08 | 0.07 | 0.16 | 0.11 | 0.04 | 0.05 | 0.02 |
| 18 | 0.00 | 0.01 | 0.02 | 0.02 | 0.01 | 0.03 | 0.02 | 0.02 | 0.02 | 0.05 | 0.03 | 0.01 | 0.01 | 0.00 |

